# PSoup: an R package for simulating biological networks from a qualitative perspective

**DOI:** 10.64898/2026.04.19.719106

**Authors:** Nicole Z. Fortuna, Brodie A. J. Lawson, Christos Mitsanis, Kevin Burrage, Christine A. Beveridge

## Abstract

Mathematical modelling is essential for understanding how complex biological systems respond to genetic, physiological, and environmental changes. Existing approaches, however, often require trade-offs between mechanistic detail, model size, parameter uncertainty, and interpretability. Ordinary differential equation (ODE) models capture biochemical processes with quantitative precision but can demand extensive parameterisation. In contrast, large statistical and machine-learning models rely on substantial datasets and frequently lack mechanistic transparency. Qualitative approaches such as Boolean networks improve scalability but may oversimplify biological behaviour. To address some of these limitations, we present PSoup, an R package that automatically converts knowledge graphs into transparent, parameter-free, qualitative models. PSoup uses algebraic update rules designed around a fixed, biologically interpretable baseline, enabling predictions of relative change across diverse perturbations without requiring kinetic parameters. This design allows PSoup to integrate information across biological scales and from heterogeneous experimental sources. We evaluated PSoup using the well-studied shoot branching network of Bertheloot et al. (2019), which incorporates hormonal (auxin, strigolactone, cytokinin) and metabolic (sucrose) regulation. Across 78 experimental conditions, PSoup correctly predicted 88.5%of perturbation outcomes, including 89.5%accuracy for unique, biologically consistent comparisons. We further demonstrate how PSoup can distinguish among alternative plausible network topologies, revealing how structural differences influence emergent system behaviour. PSoup offers an intuitive, accessible, and mathematically transparent framework for exploring biological networks. Its capacity to integrate diverse knowledge and test alternative hypotheses positions it as a powerful tool for biological discovery and a valuable complement to existing modelling approaches.

## Introduction

Mathematical modelling is an indispensable tool for understanding complex biological systems (Arrell and Terzic, 2010;Motta and Pappalardo, 2013;Nartallo-Kaluarachchi et al., 2025;Loewe et al., 2025). Modelling can be used to test whether a consolidated understanding of a system is consistent with biological reality (Craver, 2006), to make predictions of system outcomes which can be used to drive hypothesis creation, and to aid in decision making (Dun et al., 2009;Motta and Pappalardo, 2013;Brodland, 2015). Given extensive knowledge of genetics, biosynthetic pathways, and environmental responses, it becomes increasingly necessary to make use of modelling to understand how these interconnect in a causal network to dictate the behaviour of biological systems (Ji et al., 2017;Puniya et al., 2024).

There are a range of modelling approaches that are suited for different biological questions. Mechanistic models capture a bespoke understanding of biochemical or physiological processes (Gratie et al., 2013;Ji et al., 2017), whereas statistical models use algorithms to find associations in large datasets (Bzdok et al., 2019;Ghazi et al., 2022). Mechanistic models are flexible and can be used to represent a variety of interactions among diverse component types, however they can be time-consuming to build and generally limited in size. In contrast, statistical approaches, including machine learning and other large-scale modelling approaches rely on algorithmic analysis of large datasets of a common type. The trade-off is that detailed mechanistic complexity underpinning the processes of the system is replaced by complexity in the structure and scale of the network being represented (Giulini et al., 2021). Regardless of the method being discussed, there is an implicit underlying network that is being represented.

A classic mechanistic modelling approach is ordinary differential equations (ODEs). ODEs are an excellent mathematical tool to describe the kinetics that underlie biological processes. They typically use a high level of parametrisation that describes the relevant kinetics in a quantitative way (de Jong, 2002;Gratie et al., 2013;Barbuti et al., 2020), allowing for the representation of complex interactions but can restrict the number of interacting parts due to computational limitations (Fröhlich et al., 2017;Kapfer et al., 2019). A contrasting mechanistic approach is Boolean networks, which trades the parametric complexity of an ODE for an increase in network complexity. Boolean networks require defined logical rules (Chevalier et al., 2025), using binary switching through logical circuitry to determine system states (Kauffman, 1969;Thomas, 1973). This allows for increased network size while remaining tractable (Barbuti et al., 2020;Pušnik et al., 2022), and has successfully predicted phenotypes and temporal gene expression patterns (Fumiãand Martins, 2013;Davidich and Bornholdt, 2013;Grieb et al., 2015;Chevalier et al., 2025). While Boolean networks are simple in their representation of systems, they still require the user to make prescriptive decisions regarding how information is aggregated.

Statistical methods of modelling system outcomes capitalise on the informative power of complex networks and become more powerful with increasing data. Given that the algorithm is responsible for generating the network, users will not need to make as many individual decisions influencing how information flows through the network. These networks, which can consist of many thousands of nodes, are created with algorithm-based associations (Wang et al., 2021;Muzio et al., 2021;Jin et al., 2021;Becker et al., 2023). While a powerful tool for making predictions in aggregate (Lin and Zhao, 2005;Santolini and Barabási, 2018), this level of complexity is difficult to interrogate and can contain many individual errors that cancel out when considered in aggregate (Szederkényi et al., 2011;Blum et al., 2018;Stumpf, 2020). Random walks are a statistical tool that can be used to identify important nodes and pathways on a topology by stochastically exploring the network space and can also be used as a tool in machine learning methods to analyse network behaviour (Xia et al., 2020).

In general, machine learning methods generate networks implicitly in the form of black boxes. Therefore, the relevance of connections used to make predictions are difficult to assess (McCoy et al., 2022;Rudin, 2022), and the learned models are often poorly able to predict outside the range of original data upon which the model was trained (Wang et al., 2023). Graph attention networks (GAT) are a type of deep learning that can be used to find the relative importance of different connections within a graph for generating outcomes. This leads to explainable results even if the explanation does not actually match ground truth (Agarwal et al., 2023).

Researchers are increasingly making use of statistical and machine learning methods to analyse biological systems as a result of the massive amounts of data currently available. While obviously useful, we argue that there is still value in exploring known causal relationships through mechanistic modelling. The challenge in either approach is enabling connectivity across levels of abstraction (biological scales) and across disparate but rich data sources.

Many studies, grounded in a variety of modelling styles, have reinforced the idea that network structure in itself contains a high level of information regarding how a system will respond to perturbations (Kauffman, 1969;Espinosa-Soto et al., 2004;Chaves et al., 2005;Ruths et al., 2008;Fumiãand Martins, 2013;Santolini and Barabási, 2018). In addition to studies that directly test the role of the network in determining biological outcomes, the fact that a variety of parameter combinations within a given ODE model can still fit the data implicitly supports the notion of the topology itself being important for determining outcomes (Gutenkunst et al., 2007;Chis et al., 2016;Grabowski et al., 2023). These observations are also consistent with evolutionary principles in which biological systems are expected to be resilient to superficial changes (Klemm and Bornholdt, 2005;Aldana et al., 2007;Berthelot et al., 2018). We argue that focusing on network topologies to predict biological outcomes allows us to focus on the biological ‘goals’of the network and exclude detail that may be prohibitively difficult to uncover (Chevalier et al., 2025). To this end, we decided that a new modelling approach should be created that specifically generates predictions based on the network structure of the system in question.

The parent model to the method we created is that of Dun et al. (2009). This model constructed a series of 26 algorithmic equations to describe the transport of hormones between two compartments. These equations were limited to three parameters that dictated the transport of hormones between biological compartments, but otherwise did not include kinetic information. Not only did this model have a high level of success in replicating the experimental results of the system, but was also able to be used to suggest improvements in understanding. This model was explored again by Lawson et al. (2026), who not only demonstrated that the system could be expressed with fewer equations, but also showed that compared to an optimally parameterised model describing the system, there was no meaningful improvement in the capacity to predict qualitative biological outcomes of perturbative experiments. Given this conclusion, we feel justified in pursuing such a minimalist style of modelling.

In this paper, we present the PSoup R package as a method of automatically generating a simple mechanistic model from networks (knowledge graphs) built using prior scientific knowledge. Unlike traditional mechanistic models such as ODEs, we do not use parameters to represent the underlying kinetics of the system. Instead, we focus on the components in the system (nodes), the interactions between those components, and the nature of those interactions (directed and signed edges). This maintains the benefits of a model with human intelligible connections between nodes, while removing the need for representing the underlying kinetics of the system, allowing for increased complexity in terms of network size (Mitsanis et al., 2026). In addition, the flexibility in the representation of systems as knowledge graphs and by extension the model generated, means that a single model can deal with interactions across biological scales (from genetic, molecular, protein, environmental, to physical intervention).

To ensure model transparency and to accommodate diverse types of nodes in the network, PSoup uses a set of algorithmic rules to automatically translate a network diagram into a model and provides the functionality to explore the behaviour of that network through simulation. Users can design a diagram, using the freely available Newt editor (Balci et al., 2021) in the standardised diagrammatic language Systems Biology Graphical Notation –Activity Flow (SBGN-AF) (Le Novére et al., 2009;Mi et al., 2015), without being concerned with how their network is transformed into a model. This gives accessibility to modelling for biologists who may be uncomfortable with mathematics and makes the process of updating models straightforward.

Our algebraic rules are designed such that the system’s equilibrium state, absent any perturbations, is a value of one (1) at all nodes. This defines the baseline as wild-type or control, and then all model predictions under different scenarios can be naturally interpreted relative to the baseline. Defining model outputs in this relative way has the benefit that the network topology can be updated without changing the outcome of the baseline condition. Additionally, we demonstrate how our update rules can be derived from standard ODE approaches for systems biology under certain assumptions, hence providing a principled means of customising these rules where those assumptions might be deemed inappropriate, even for this qualitative modelling context.

### Design and Implementation

PSoup is an R package that translates knowledge networks into algebraic formulas, by following a set of algebraic rules. Included in the package are a series of tools to explore the behaviour of networks in response to perturbations. To find tutorials on how to use PSoup see https://nicolezfortuna.github.io/PSoup/.

### Features of a PSoup model

A PSoup model begins with a knowledge graph created using a subset of the SBGN-AF diagrammatic language (Figure 1). The knowledge graph contains meaningful components of a biological system and how they interact, potentially across multiple biological scales, from molecular through to organs or environment. The structure of the knowledge graph is provided as input in the form of an .sbgn file to the PSoup package, which generates the update rules that allow the modeller to simulate the system and how it responds to perturbations.

**Figure 1.**
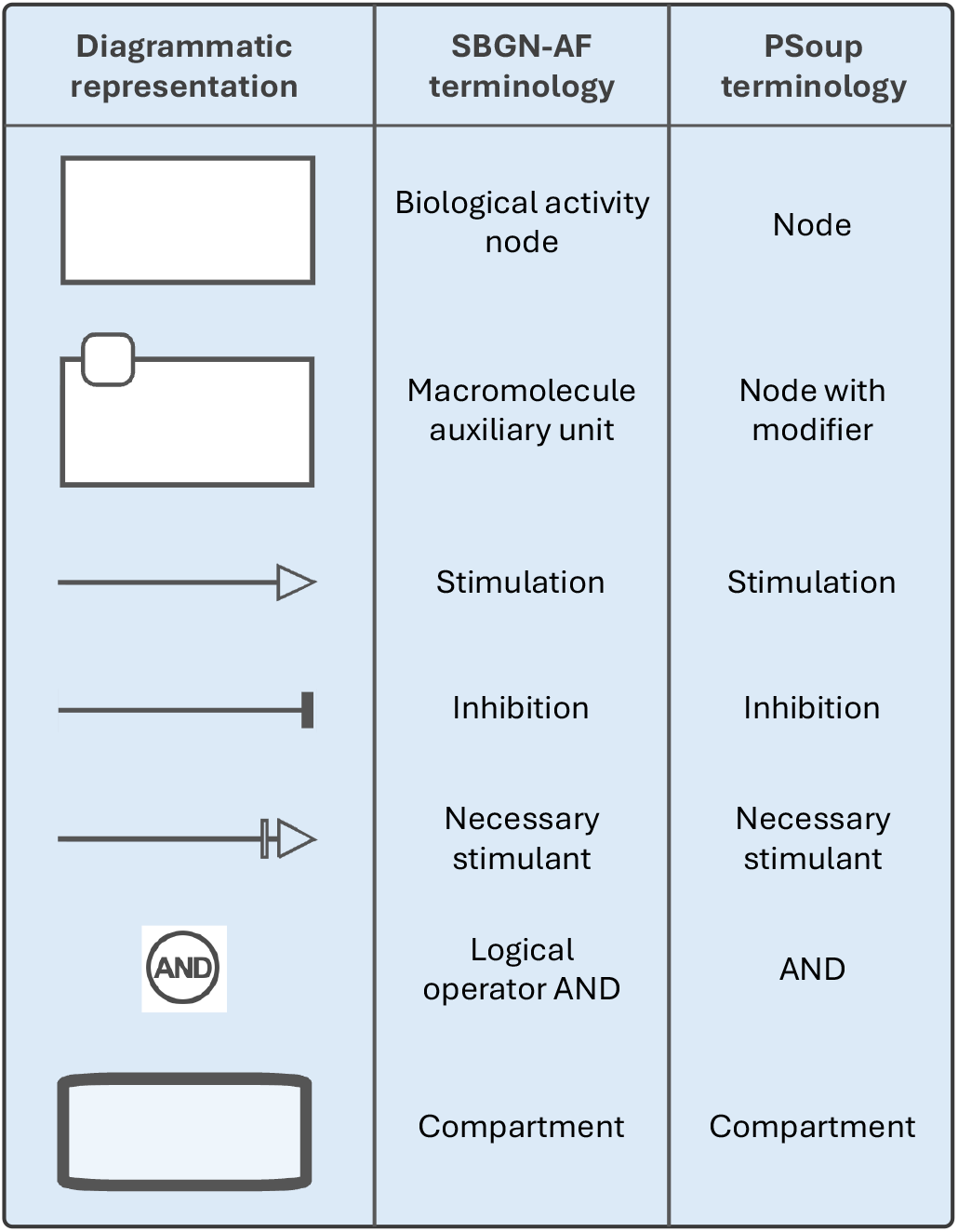
A list of all the symbols that can be used to construct a PSoup model, including the original terminology from the SBGN-AF diagrammatic language.

There are four units of construction that can be used to design a knowledge graph for PSoup, all which can be represented using a subset of the SBGN-AF language as seen in Figure 1. The first are the network nodes that represent the components of the system across multiple levels of hierarchy. Within the context of a biological system they can represent genes, hormones, signalling molecules, enzymes, environmental signals, and transcription factors. Nodes can also be used to abstractly represent phenotypic traits of interest, such as branching, height, or fruit quality.

The second unit of construction are the edges that connect nodes and dictate the effects that upstream nodes have on downstream nodes. The SBGN-AF networks used to generate a PSoup model are signed and directed, meaning that edges indicate the direction and the type of effect between nodes. There are three edge types that a PSoup model recognises: stimulation, inhibition, and necessary stimulation. The standard stimulatory and inhibitory edges indicate whether the upstream node has a positive or negative impact on the downstream node respectively. Necessary stimulants are distinguished from ordinary stimulatory edges by the specific property that if any of them are removed, the downstream product will be eliminated regardless of the action of any other nodes that regulate it.

In addition to the three edge types, edges can be augmented through the operator AND, in cases where the influence of upstream nodes are dependent on each other. An example of when AND might be useful, is when an intermediary complex must be produced, or if a product must interact with an enzyme to induce the downstream effect. The behaviour of an AND operation in PSoup requires all of its inputs to be present in order to pass along its effect.

The third unit of construction, modifiers, are a way to indicate where perturbations can occur in the network. Modifiers are attached to nodes to indicate that the node can be weighted to achieve a network perturbation. For example, if a node represents the action of a hormone, genes that are known to influence the production of the hormone can be listed in the modifier. Through this modifier, users will be able to knock out, reduce the expression, or over-express that node.

As the fourth unit of construction, networks can be partitioned into compartments. This can be used to model tissue specificity in multicellular organisms, or even to represent grafting experiments. In plant science, grafting can be used to combine different genotypes into a single plant. PSoup can represent such experiments as it permits nodes and modifiers of the same name to be used in different compartments, with the potential for perturbations to be differentially applied between compartments.

### Diagrammatic language behind PSoup

The choice of the SBGN-AF language enabled us to represent all four units of construction required for a functional PSoup model in a knowledge graph, enabling the automatic generation of the model. Diagrams using SBGN-AF notation can be built using a freely available online tool called Newt Editor (https://newteditor.org/). The subset of SBGN-AF diagrammatic notation used to generate a PSoup model can be seen in Figure 1.

### Rules for generating a PSoup model and its usage

Once a knowledge graph has been created with the Newt Editor, PSoup uses algorithmic rules to convert it into a series of equations. The most important design principle of PSoup is the notion of representing systems relative to a baseline value of one (1), making predictions unitless and qualitative. As mentioned above, we specifically designed the set of algebraic rules so that the baseline condition for any PSoup network model produces a steady state value of one for all nodes. Perturbations are introduced to examine direction of change across the network relative to the baseline.

Perturbations are specified using the modifiers attached to network nodes, which are normalised such that a value of 1 represents the unperturbed state. A modifier’s value can be set to a value <1 to represent downregulation of that node, and similarly values >1 represent upregulation. A modifier set to 0 fixes a node’s value at 0, representing removal of that component from the system (for example, representing a gene knockout experiment). Once a system is perturbed through modifiers, node values >1 represent increased response, while values between 0 and 1 represent a reduction in response.

### Constructing equations

PSoup models are difference equations that describe how a system’s state evolves dynamically over time in discrete steps, using an iterative application of an update rule. Usually, the steady state attained by the model is the property of interest, which can be obtained by simply performing many such iterations. However, as we detail in Supplementary Five, the PSoup update rules correspond to numerical integration of a specific ODE model and hence represent an approximation to genuine biochemical dynamics.

The general form defining the rule that updates node values in PSoup consists of the product of factors corresponding to the different types of influence that PSoup networks include (Equation 1). The values of a node’s modifiers also appear in this equation. These behave as input variables in the model and scale the value of nodes independent of the effects of network interactions.

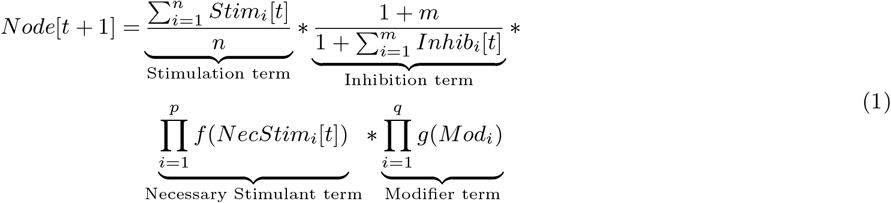

Stimulants and inhibitors are represented using simple mathematical forms that satisfy the fundamental properties of increasing with the values of stimulus nodes and decreasing with the values of inhibitor nodes. To achieve a fixed baseline that is robust to the addition or removal of nodes and/or edges to a network’s structure, the effects of incoming stimulatory and inhibitory edges need to be normalised against the number of incoming nodes of each type. These considerations motivate the choice to take the mean value of incoming stimulants or inhibitors for each respective term. The values of all nodes are updated simultaneously.

*Stim*_*i*_ and *Inhib*_*i*_ denote the *i*-th stimulant and *i*-th inhibitor for the node of interest. *n* and *m* denote the numbers of stimulants and inhibitors for that node, respectively. Biologically, these terms imply that each individual stimulant or inhibitor for a node is equally important and that the more stimulants (or inhibitors) a node has, the less each contributes individually to its value. This is a natural consequence of the choice that stimulants and inhibitors act independently, and that the baseline should be a fixed value.

For necessary stimulants and node modifiers, a product of all incoming nodes of the same type is used. *NecStim*_*i*_, and *Mod*_*i*_, give the *i*-th necessary stimulant and *i*-th modifier for the node of interest, with *p*, and *q* denoting the numbers of each. The user has the flexibility to specify the function (*f* and *g*) that adjusts the precision with which the values of the necessary stimulant nodes and modifiers contribute. These functions are the identity function by default, *f* (*x*) = *x*, but any function satisfying *f* (0) = 0 and *f* (1) = 1 is a valid choice, and we discuss alternative forms representing different kinds of biology in Supplementary Six. Using a product of necessary stimulants (and similarly for modifiers) ensures that if any of them takes a value of zero then the value of the node at the next timestep is also zero. This captures the scenario where multiple components are required for synthesis, and there is no product if any are missing.

In addition to the general form, the operator AND can be used to modify the way that upstream nodes influence the downstream node (Equation 2). This is useful if two or more upstream nodes are non-independent in their influence on a downstream node (for example, if an intermediary complex must be formed, or enzymatic action must take place). Upstream nodes that are part of an AND are treated as a single input in the general form, with a value that is taken from the geometric mean of the nodes involved in the conjunction. The form chosen for an AND ensures that despite any number of non-independent effects on a node, the influence of the AND remains at the baseline of 1 if all members of the conjunction have the value 1. It also results in a value of 0 if any input into the AND is 0.

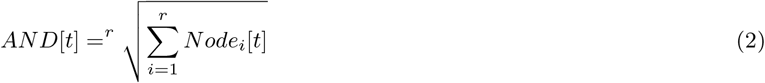

*Node*_*i*_ represents the *i*-th node involved in the AND operation, the final value of which will operate as if it is a single node within the PSoup propagation algorithm (Equation 1).

Compartments can be useful in the generation of equations for cases of tissue-specific gene expression or multiple sites of production of the same product. They allow for the same name for nodes and modifiers to be applied across different compartments. PSoup automates the independence of the nodes by producing compartment specific equations for nodes that exist within multiple compartments.

#### Dependencies

The PSoup R package depends on several supporting packages: xml2 (≥version 1.3.3), ggplot2, methods, stats, utils, prodlim (≥version 2024.06.25), stringr, reshape2, parallel, parallelly, foreach (≥version 1.5.2), and doParallel (≥version 1.0.17). It was built in R version 4.1. PSoup requires diagrams in the SBGN-AF language as input.

#### Model validation

One design benefit of the PSoup approach is that knowledge graphs can be created from diverse sources of information. The challenge then is to validate the PSoup predictions against these diverse sources. When validating a PSoup prediction, a baseline condition (usually the control or wild-type) will be chosen for each set of comparable experiments used for developing the knowledge graph. Once the baseline condition is identified, it is used to normalise experimental conditions such that they can be used as inputs for PSoup simulations. For example, a knockout mutant would mean the associated modifier representing the relevant gene would be given an input value of zero. The outputs of experiments are also normalised into categories of greater or less than the baseline output. These normalised experimental categories can then be compared with the output of PSoup.

Once experimental results and their simulated counterparts have been compared, mismatches should be inspected and critically evaluated. If an error has been found in the diagram, the knowledge graph should be rebuilt and the process restarted (Figure 2). In some rare cases, the mismatch could be due to an ambiguity regarding the experimental biological data. To resolve possible ambiguities, two checks should be made. First, is whether there are any repeated instances of that experimental condition and whether these repeats are biologically consistent. Second, it should be checked if the biological difference was statistically different. If the mismatch was not biologically significant, as judged from publications as overlapping confidence intervals, the condition should be flagged for removal from downstream analysis. If the remaining condition sets are biologically significant and yet still inconsistent, that condition set should be removed as there is no biological consensus on the expected outcome of that perturbation. Such an occurrence reveals a limitation in our biological knowledge.

**Figure 2.**
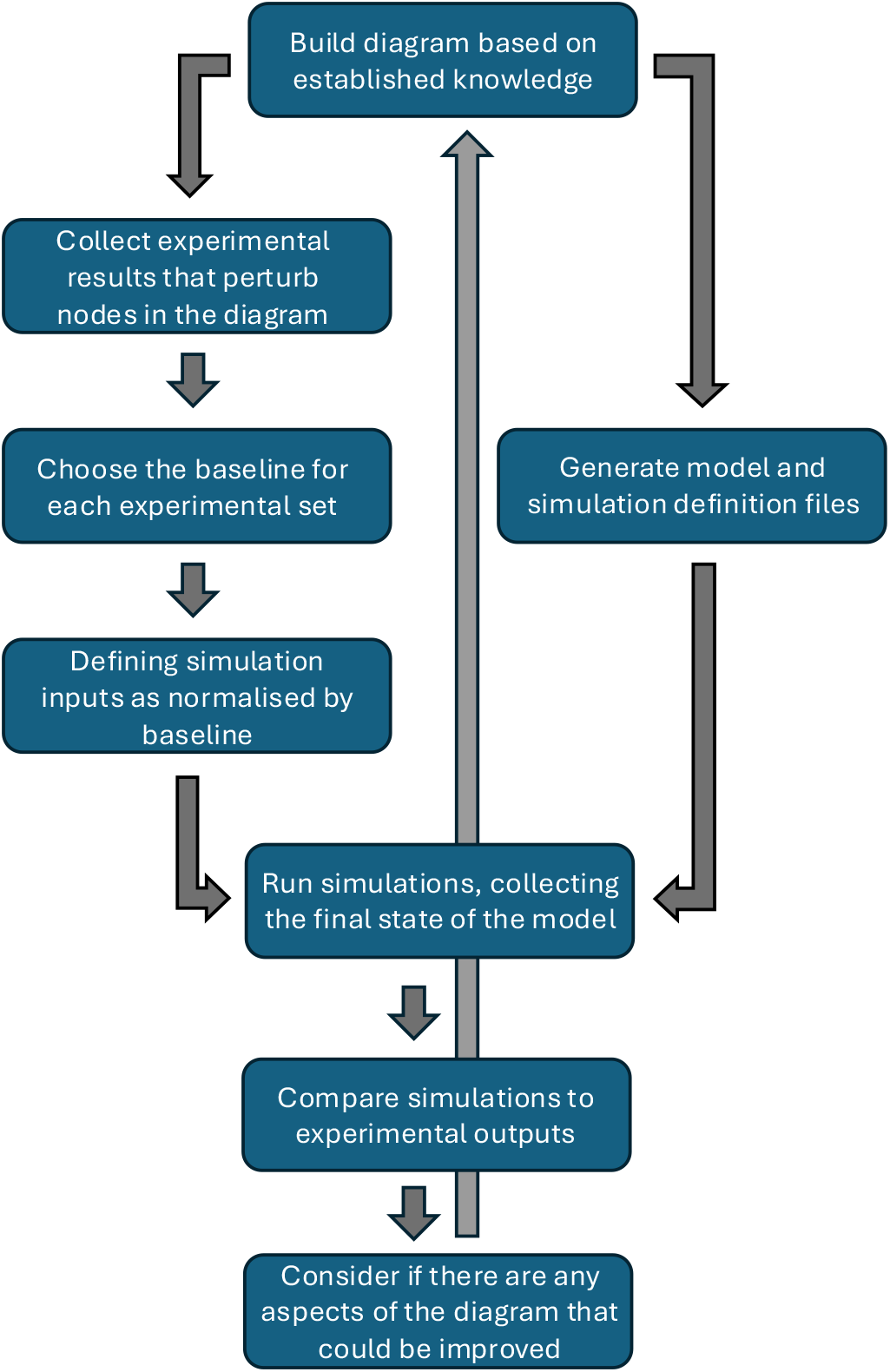
A flow chart of the progression for building and analysing a PSoup diagram.

#### Shoot branching Case Study

The main purpose of developing this PSoup method was to determine whether current understanding of biological processes (knowledge graphs) can be validated using a modelling approach that is standardised and has not been highly engineered by extensive parameter fitting. As such the PSoup method needs to be applicable for a plethora of experimental approaches such as is typical for plant physiology and development. As a demonstration of the functioning of the PSoup approach, the Bertheloot et al. (2019) model of branching was chosen (henceforth referred to as the Bertheloot model).

The Bertheloot model is simplistic in terms of the number of nodes (6) included, yet highly complex in that it was created with a series of ODEs that represent a total of 19 parameters. This ODE model was validated using a set of specific phenotypic outcomes that the experiments themselves did not always measure. Therefore, to validate the model, some of the experimental outcomes needed to be used as the input for a separate equation which calculated the values used in their ODE model. This means that the approach does not enable direct testing across the complex range of wet-lab experiments conducted. We hypothesised that the PSoup approach can be used to validate against each and all relevant experiments, ensuring that the knowledge graph is sufficient to explain various data types.

The Bertheloot et al. (2019) study is an ideal choice for identifying whether the PSoup approach can achieve such a broad validation assessment because experiments included (i) two plant systems: rose and garden pea, (ii) heterogeneous conditions across two different laboratories;(iii) plants in pots and studies of isolated buds in vitro;and (iv) diverse treatments such as defoliation, decapitation, hormones, sucrose, and wild-type and mutant plants. It is therefore a great example study to test whether the PSoup method is well suited to evaluate and test networks purporting to represent knowledge from diverse experimentation.

The Bertheloot model is a mathematical model of the simplified processes controlling shoot branching, whereby the plant hormone auxin inhibits branching by positively and negatively regulating two other hormones, strigolactone and cytokinin, respectively. In turn, strigolactone inhibits, while cytokinin promotes shoot branching. At the same time, sucrose, the main product of photosynthesis, suppresses the action of strigolactones and promotes cytokinins to lead to the promotion of branching (Barbier et al., 2019;Beveridge et al., 2023). The study includes a series of experiments primarily focussed on testing the role of sucrose relative to hormones in controlling shoot branching. The study focuses on how various single and multiple interventions work within a network to control branching (and occasionally, levels of the hormone cytokinin).

#### Bayesian Inference for the Bertheloot Model

In order to make our case study more complete, and demonstrate the difficulty of using typical biological data to estimate the values of parameters in quantitative models, we perform Bayesian parameter inference (Linden et al., 2022) for the Bertheloot Model. This inference functions similarly to the calibration used to select parameter values in their study and makes use of the same data, but the Bayesian approach also produces quantitative estimates of the uncertainty that remains in these parameters after taking into account the data. Samples output by the inference routine represent different data-consistent calibrations of the model, and comparing these calibrations indicates what the data do or do not consistently suggest to be true about the model’s behaviour.

Specifically, Bayesian inference quantifies parameter uncertainty by describing beliefs about a model’s parameters, *θ*, in terms of probability distributions. Bayes’theorem links these beliefs prior to considering the data, *p*(*θ*), to the beliefs after seeing the data, *p*(*θ*|*d*),

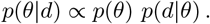

This is done by using the model (together with some specification of error) to define the likelihood, *p*(*d*|*θ*), that the data would be observed given some value for the parameters. In general, the normalising constant (the constant of proportionality) is unknown, and inference is performed by generating samples from *p*(*θ*|*d*) using methods that do not require the normalising constant to be known (Kroese et al., 2011).

As quantitative models and Bayesian inference are not points of focus in our study, we reserve full discussion of the inference process for the Supplement (see Supplementary Five). To summarise the process, however, we first make a minor re-parameterisation of the model such that its parameters have clearer physical interpretations, then apply a log-uniform prior (uniform in scale) to all parameters, reflecting no solid beliefs about the biochemical kinetics of shoot branching before seeing the data. For simplicity, we choose a Gaussian error model, as is common (Lambert et al., 2023), and this together with the observed data defines the likelihood. The data are presented in Table S4 of the Bertheloot et al. (2019) study’s supplementary material, consisting of different levels of synthetic auxin and sucrose that were supplied to plants as experimental treatments, and the associated measurements for bud outgrowth time and plant levels of strigolactone and cytokinin. Samples from the posterior are generated using the resample-move variant of sequential Monte Carlo (Gilks and Berzuini, 2001), an approach that is highly parallelisable and that helps avoid poor mixing on difficult posteriors by gradually introducing the difficulty of the sampling problem through likelihood annealing (Neal, 2001). We use the adaptive selection of the number of move steps presented by Drovandi and Pettitt (2011).

## Results

### Multiple parameter combinations match biological data

PSoup uses a qualitative, but biochemistry-informed description of biological networks that avoids the need for parameter calibration. Our branch inhibition case study serves as a good example as to why this type of modelling approach is important, even when quantitative data are available. As part of their study, Bertheloot et al. (2019) manipulated cytokinin and strigolactone levels and recorded the time until bud outgrowth (or failure to bud) for twelve different experimental scenarios where different amounts of auxin and sucrose were supplied to axillary buds. These and other quantitative measurements, such as cytokinin levels were used to calibrate a biochemical kinetic model that predicts the same three measured properties, given a level of auxin and sucrose.

Such data can certainly be used to calibrate this type of quantitative model, however simply choosing the best-fit values for the parameters does not properly consider whether the data are sufficient to rule out other possible calibrations. For models in systems biology, the existence of many possible fits is if anything the expected scenario (Gutenkunst et al., 2007). Here, by taking a Bayesian inference approach, we instead obtain a probabilistic description of the whole space of plausible calibrations for the model. This allows us to consider which model predictions are consistently supported by the data, and which are specific to some certain choice of parameter values. As Figure 3 demonstrates, even between models that agree with all the data, the quantitative predictions can still differ significantly.

**Figure 3.**
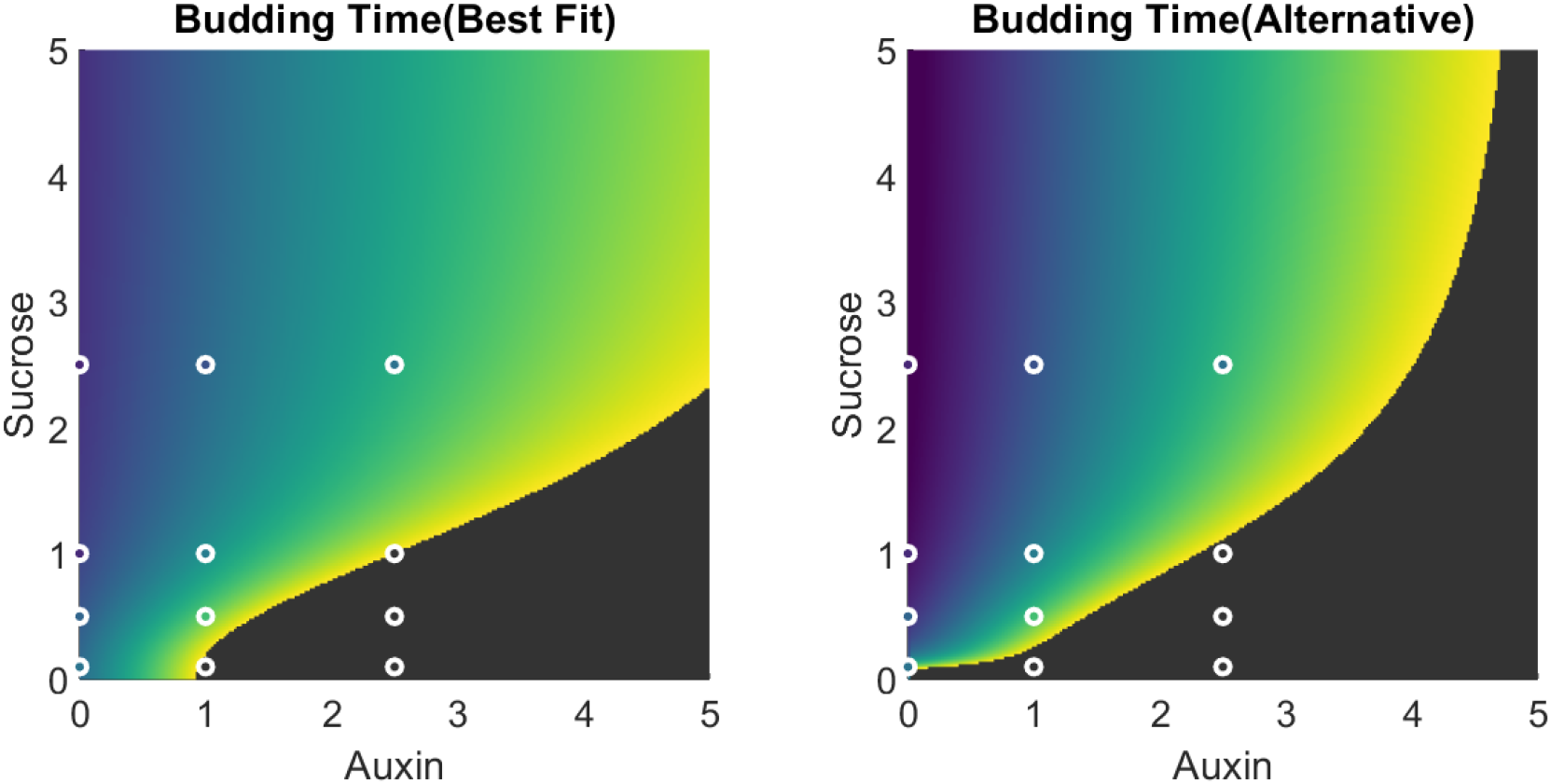
Similarly plausible calibrations to quantitative experimental data can produce models that make very different predictions. Heat maps show the budding time (in days) for an axillary bud when supplied with different levels of auxin and sucrose. Dark regions indicate a complete failure to bud. The colour inside white circles shows the experimentally observed budding time under those auxin and sucrose levels. In particular, the models’predictions about failure to bud when auxin and sucrose are both at a low level or both at a high level are very different.

Consistently, the effect of sucrose is to shift budding time earlier (or a bud closer to release), and auxin has the opposite effect. However, the manner in which auxin and sucrose interact (through their downstream targets cytokinin and strigolactone) to modulate bud outgrowth differs across the set of data-consistent models. In the example shown in the left panel of Figure 3, the best-fit (maximum a posteriori, in Bayesian inference terms) model suggests a comparatively weak effect of auxin, with a moderate supply of sucrose sufficient to activate bud outgrowth even when auxin supply is doubled beyond the highest level experimentally tested. However, the same chemical kinetics can also be fit to the data (Figure 3, right panel) in such a way that suggest that high auxin levels have strong enough effect that none of the sucrose levels considered are able to induce budding. Additionally, when the supplied sucrose is low, the alternative model predicts that a much smaller amount of auxin is required to switch off budding.

Furthermore, the Bertheloot et al. (2019) biochemical kinetic model (laid out in Supplementary Five) is itself a simplification of the true biochemistry. Standard mathematical forms for describing activation and inhibition were used, with arbitrarily-chosen Hill coefficients. Aspects of the kinetics, such as the way in which sucrose acts to hamper strigolactone perception, were treated phenomenologically. We do not raise these issues to criticise the Bertheloot et al. model, but rather to highlight that even if much more data were available, the predictions of the model could still very well depend on the modelling choices used to construct it.

The PSoup approach rather accepts that the true biochemical kinetics of a system of interest are unlikely to be fully known, and that the data required for rigorous, uncertainty-free calibration are rarely available. Instead, the basic character of a network’s implied biokinetics is extracted and the resultant model simulated under different scenarios to compare its predictions in a qualitative fashion. As we demonstrate using the branching inhibition case study in the following sections, this type of modelling still allows for a rich investigation of the biology of a system, including the testing of alternative hypotheses.

### PSoup has a high success rate of capturing biological outcomes

The Bertheloot model of branching was redrawn (Figure 4) using the PSoup subset of the SBGN-AF diagrammatic language. This figure was then used as an input to PSoup to produce the mathematical model to be used to simulate the system (Equations 3 to 8). In the SBGN-AF network (Figure 4) four nodes (*Auxin, Strigolactone, Sucrose*, and *Cytokinin*) contained modifiers (*Aux, SL, Suc*, and *CK* respectively) which were used to indicate points in the network that could be perturbed. One node was used to integrate information (*Integrator*), and one node was used to track the phenotypic output (*Branching*). The word integrator is used to capture the process by which sucrose affects strigolactone action, not to be confused with the transcription factor BRC1 (Aguilar-Martínez et al., 2007). The *Auxin* and *Sucrose* nodes act as feeders as they have no upstream influences, and the *Branching* node as a terminus since it does not exert influence on any other node. The branching node in our network was typically used as the predicted value in our simulations and was used to compare with the various branching outcomes measured in the Bertheloot et al. (2019) study. However, any node can be used for prediction given perturbation at another node(s). In fact, *Cytokinin* was occasionally used as a phenotypic outcome.

**Figure 4.**
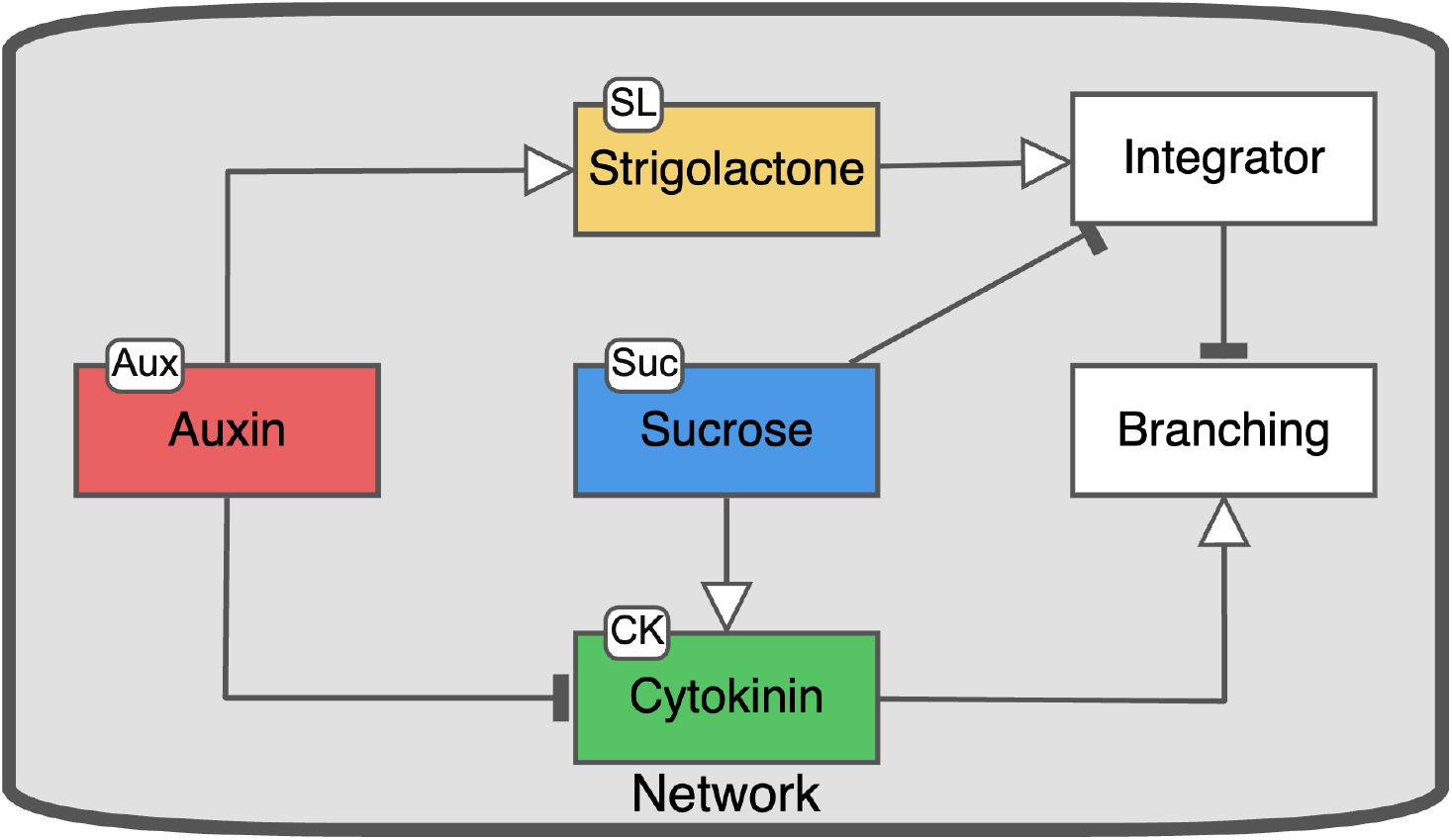
A depiction of the branching network as described by Bertheloot et al. (2019), created using PSoup compatible SBGN syntax within the Newt Editor software.

The model generated by PSoup based on the redrawn Bertheloot model resulted in a set of 6 difference equations. The terms in the equations below match those found in Figure 4:

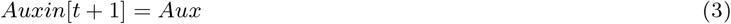

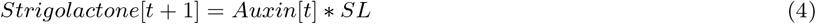

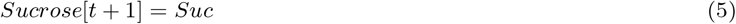

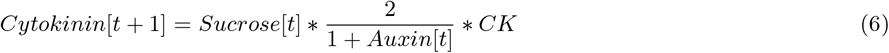

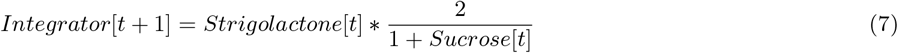

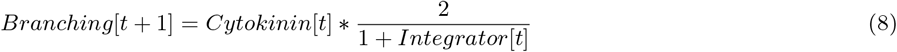

As a result of *Auxin* and *Sucrose* both being feeder nodes in this model, both functionally behave as constants. In addition, given that the maximum distance that information can travel is only three steps, this model always resolves to its equilibrium state quickly. For a more complex model, especially one that contains feedback loops, there may need to be many iterations of the propagation algorithm to find the steady state of the system.

The experimental wet-lab results of the paper were summarised in a spreadsheet, organised by figure of origin, noting the conditions that would affect each node for each experiment (Supplementary One). For each experiment, one condition was chosen to represent the baseline control against which other conditions were compared. In some experiments, wild-type plants were not included and hence an alternative baseline was chosen. Supplementary One contains the full summary of the biological data, including the chosen baseline for each Bertheloot figure, and the reasoning behind the removal of the few experimental conditions excluded from this study. The information in Supplementary One was used to produce Supplementary Two, which redefines the experimental conditions relative to each baseline so that they could be used to define simulations.

The experiments produced several different branching measures. Branch number or length both indicate more branching with increasing values. In contrast, the time to bud outgrowth or time to achieve a particular branch length indicates less branching with increasing values. Occasionally, experiments also measured the level of cytokinin in the system. PSoup simulations were run until the model settled on a steady state, with the final values defining in what direction the system was pushed in response to perturbations. Finally, a comparison was made between the simulated outcomes and the experimental outcomes (Figure 5). The simulations predicted an abstract ‘branching phenotype’relative to a baseline that needed to be compared with various experimental phenotypes related to branching. A condition was said to be validated if the simulation and the experiment moved in the same direction in response to perturbation.

**Figure 5.**
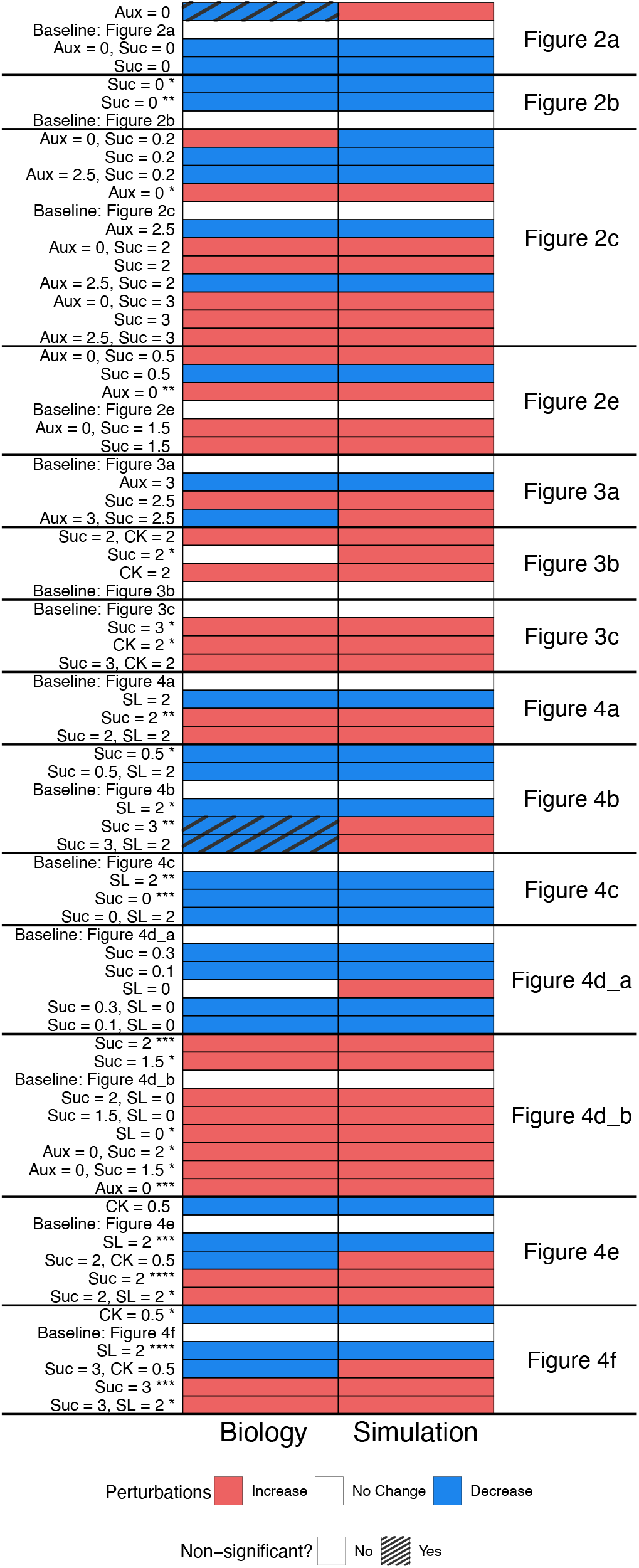
A comparison of the experimental results from Bertheloot et al. (2019) (left), with the simulated results generated using PSoup (right). Data are organised according to the figure of origin within Bertheloot et al. (2019) and shown relative to a designated baseline (control) for each experiment. In instances where the biology and the simulations did not match, cross hatching was used to indicate if the biological outcome was not significant (had overlapping confidence intervals with the baseline condition). To indicate if a condition has already been represented, the *symbol was used to show the number of repeats.

Figure 5 depicts the full comparison between the experimental results of the Bertheloot paper (left) and their simulated recreation in PSoup (right). Included conditions are organised according to the figure of origin for the data, and the modifiers used to define the simulation are given for each condition. The full biological description of the experimental conditions can be seen in Supplementary One. For conditions where there wasn’t agreement between the biological outcome and the simulation, the original biological outcome was checked to see if it was statistically different from the baseline condition for that experiment. Those that were not significantly different were denoted with cross hatching to indicate that there is not strong support for that biological result. 88.5%of the simulations matched the outcome of the experiment (69 out the 78 included perturbations). However, several of the perturbations relative to baseline were repeated between experiments. Every time a perturbation was repeated, it was indicated with a *after the label, with the number of *indicating the number of times that condition had been repeated.

### Making unique comparisons

Of the tested conditions 22 were completely unique, and 15 had between 2 and 5 repeats consisting of studies that addressed the same comparisons, although in a variety of these different systems. Figure 6 represents only the outcomes of unique comparisons. All baseline conditions have been removed as by definition they will create a biologically valid result and therefore cannot be used to test the validity of the model. Any of the perturbation types that had been represented more than once were assessed for biological consistency (that they all moved in the same direction in response to the perturbation across experiments). If the biology was inconsistent, this condition type was removed as it cannot be used to test the validity of the simulated outcome. The exception to this is if the one inconsistent biological result was found to lack statistical support in the original biological experiment.

**Figure 6.**
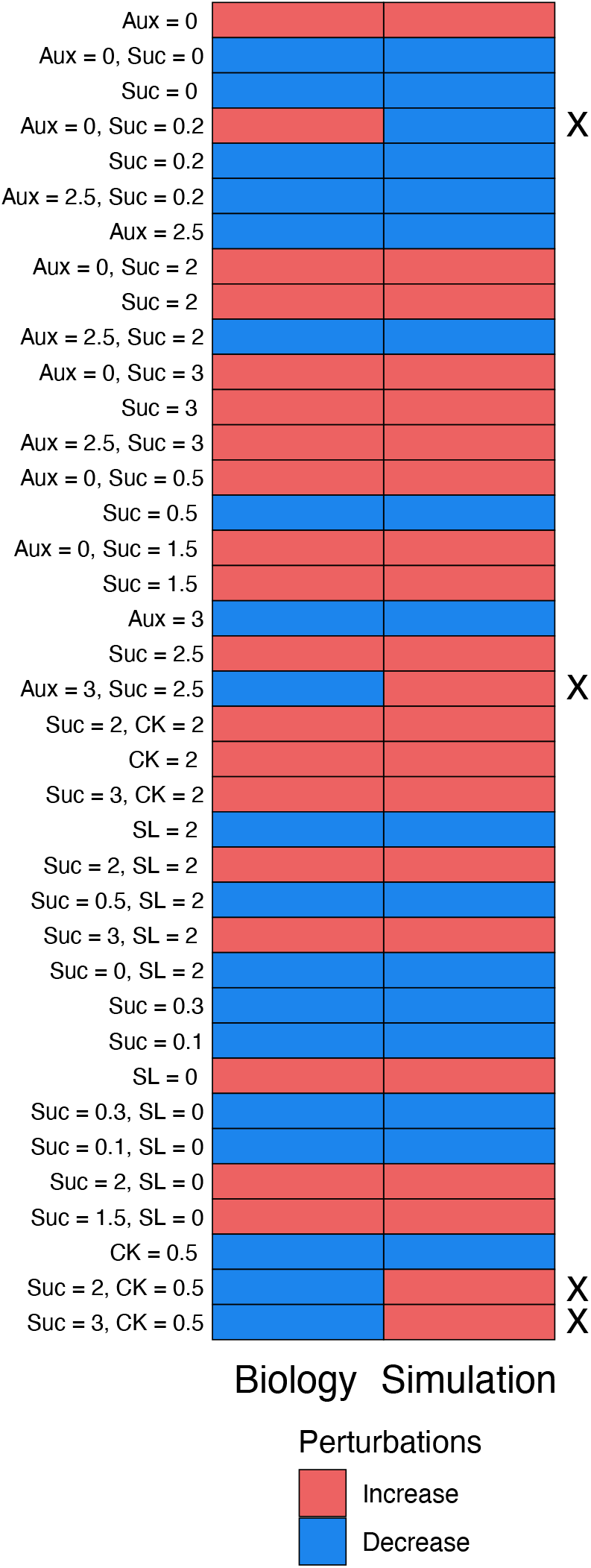
A comparison of the unique consistent experimental results from Figure 5 (left) and corresponding simulated outcomes showing the direction of change in the measured phenotype relative to the baseline condition (right). The columns show the effect of the network perturbation as either an increase or a decrease in the measured trait. An x has been used to show any instances of a mismatch between the biological and simulated result.

Of the unique conditions included in Figure 6, 89.5%(34 out of 38) matched between the experimental outcomes and the simulations. In the following repeated sets of conditions: *Aux* = 0, *Suc* = 2, *Suc* = 3, and *Suc* = 3 and *Sl* = 2, one of the repeats within each set was inconsistent. In each case, the mismatch was not statistically different from the baseline condition in the original biological experiment, and so the mismatch was removed, and the condition kept. The PSoup method had a very high level of success for predicting the direction of perturbation in the system described by Bertheloot et al. (2019). All the pairwise comparisons (conditions that had a single point of change) predicted the correct category. The only mismatches were for conditions where two perturbations of opposite effect occurred together. Errors in such a case is natural given the lack of kinetic parameters in the model, given that if one influence has a greater contribution to the phenotype than another, PSoup has no mechanism to reflect these differing contributions. Even so, of the double perturbation cases (12), 8 predicted correct results (67%).

In developmental genetics and physiological studies, comparisons for double interventions are made with each of the individual interventions. Therefore, we made sure that PSoup was able to capture the order of phenotypic outputs given different combinations of interventions. In all cases, PSoup was able to capture the direction of such trends, with minor caveats. See Supplementary Three for a full comparison of the simulated data, compared with the normalised biological outcomes.

### Exploring alternate networks

A central purpose of PSoup is to provide a robust method to test collective understanding from diverse studies through a method of qualitative prediction. It is hoped that the framework can be used to identify and evaluate alternative hypotheses. To produce an example of this, we explored three neighboring networks, each with some edge difference from the Bertheloot network, that represent reasonable minor modifications. These neighboring networks were explored to test the robustness of the method to reasonable biological interpretation, as well as to see if any alternatives were better able to explain the biological data. This exploration shows how readily the approach can shift research emphasis from estimating parameters towards thinking about alternative network structures.

In Figure 7, the original Bertheloot network topology (Fig 7Ai, reading from left to right) can be seen along with three alternatives. The first alternative (Fig 7Aii) relates to removing the stimulatory edge from sucrose to cytokinin. While there is evidence for this connection within the literature (Barbier et al., 2015;Kiba et al., 2019), the evidence for this being a strong connection appears weak given the results within the Bertheloot paper itself, which suggested only a minor contribution from cytokinin towards the final branching phenotype. Given that the model we are exploring is only representing the most important relationships in the branching network, we found it reasonable that someone might choose to omit this connection altogether if they were basing their network solely on the evidence found within Bertheloot et al. (2019). The second alternative topology introduces an additional stimulatory edge between sucrose and the branching outcome (Fig 7Aiii). This edge reflects that there are likely alternative routes by which sucrose influences the branching phenotype other than via strigolactone and cytokinin (Fichtner et al., 2021b). The third alternative topology both removes the stimulatory edge between sucrose and cytokinin and adds a stimulatory edge between sucrose and the branching outcome (Fig 7Aiv).

**Figure 7.**
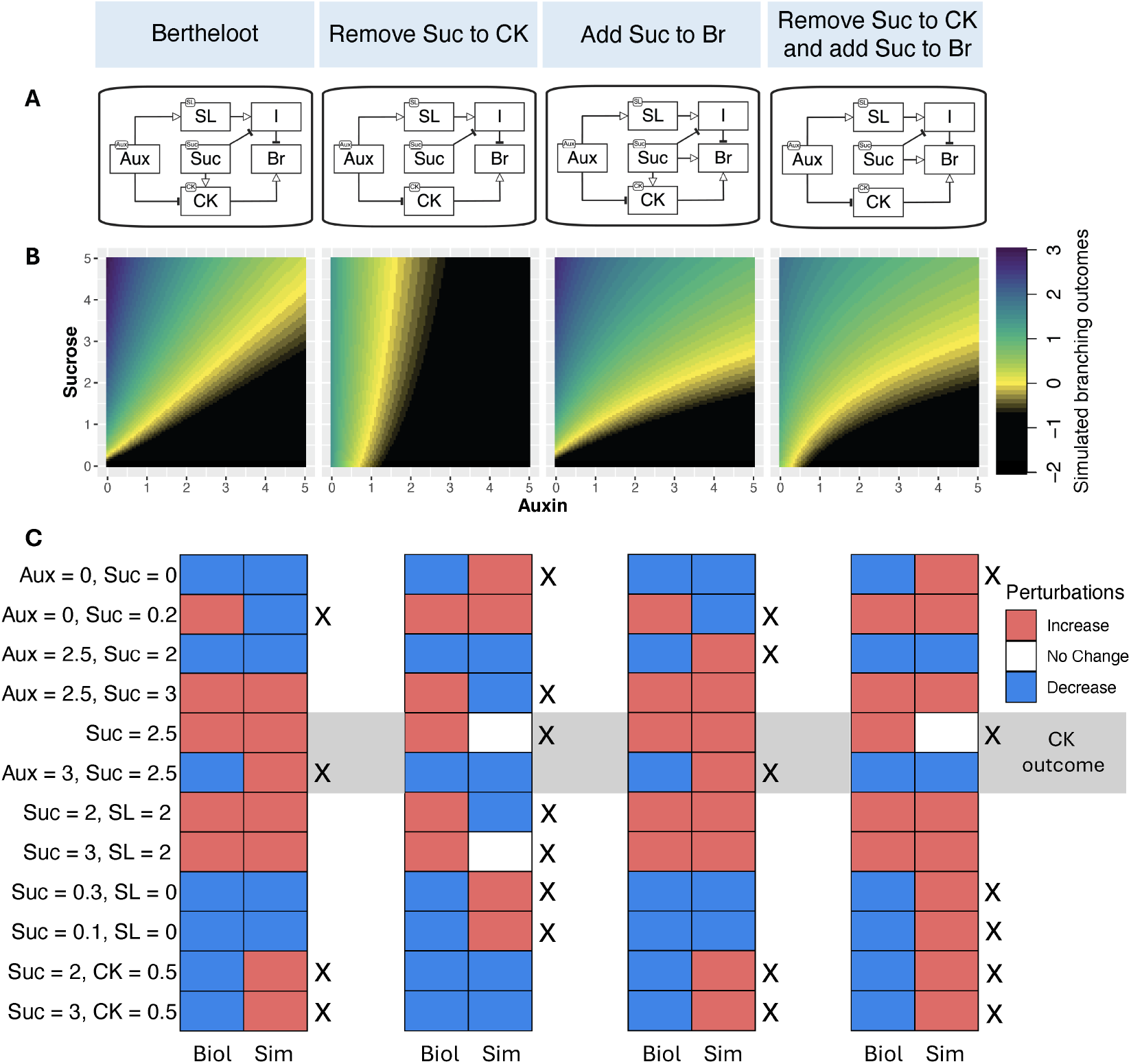
A comparison of the performance of three alternative networks against the original Bertheloot model. A) The four different topologies tested. B) The respective branching landscape when two variables, Auxin and Sucrose, are explored. In these figures, the log of the branching outcome has been shown, so that the value of 0 represents the baseline condition. Note: the scales in the x and y axes of the landscapes bare no relation to that found in Figure 3. C) Comparisons of the performance of each individual model when compared to the biological outcomes in the Bertheloot 2019 paper. Only conditions that showed a mismatch in at least one of the networks was included. Xs have been used to draw attention towards mismatches for ease of comparison. For conditions where the phenotype being tested was cytokinin rather than the branching outcome, a grey background has been used.

To understand how these topological changes (Figure 7A) modify the behaviour of the system, we tested how they responded to identical perturbations. In Figure 7B, the four networks were used to produce a series of different branching landscapes in response to perturbations to the sucrose and auxin modifiers. The Bertheloot landscape (Fig 7Bi) shows a fan pattern, whereas the other models have non-linear dynamics. The removal of the sucrose to cytokinin stimulatory connection caused the most extreme dynamics (Fig 7Bii). The two topologies that included a stimulatory connection between sucrose and the branching outcome produce very similar landscapes, with the primary difference between the two being the outcome when both sucrose and auxin are close to 0 (Fig 7Biii and 7Biv). Figure 7C is a comparison of the outcomes relative to the experimental biological conditions explored in Bertheloot et al. (2019). Only conditions that produced incorrect results in any of the models were included, to highlight conflicting results between the models (to see the full comparison of unique conditions for each alternate model, see Supplementary Seven). These represented 12 of the 38 unique conditions, with the highest number of conflicting results for an individual model being seven (81.6%accuracy). Almost all of these 12 conditions represent instances containing two opposing perturbations. This demonstrates that there was a reasonable capacity for each knowledge graph to capture biological outcomes.

The best performing model was that based on the original Bertheloot topology with only 4 incorrect predictions out of 38 (89.5%accuracy). The second-best model was when an additional stimulatory edge was added between sucrose and the branching outcome. This model produced predictions almost identical to that of the Bertheloot topology, with one additional incorrect prediction when Aux = 2.5 and Suc = 2 (86.8%accuracy). The third-best model maintained the additional stimulatory edge between sucrose and the branching outcome but removed the original stimulatory edge between sucrose and cytokinin. This model had a total of 6 incorrect predictions (84.2%accuracy), only two of which overlapped with the Bertheloot model. The worst model, which completely omitted alternate routes of action for sucrose outside of strigolactone, had 7 incorrect predictions (81.6%accuracy). This outcome aligns with the known fact that sucrose has multiple routes by which to influence branching phenotypes (Salam et al., 2021;Fichtner et al., 2021a;Beveridge et al., 2023). Interestingly, the mismatches of this model have no overlap with the mismatches of the original Bertheloot model.

For the alternative network where an additional stimulatory edge is added between sucrose and the branching outcome (Fig 7Aiii), as is the case for the original Bertheloot model, if a physiological/developmental approach of comparison is made where double perturbations are compared with their single counterparts, then the biological and simulated outcomes follow expected trends. For the two remaining networks (Fig 7Aii and 7Aiv) which represent cases where the stimulatory connection between sucrose and cytokinin has been removed, each includes a particular mismatch. Specifically, the only ‘single’perturbation shown, where the effect of sucrose alone on cytokinin is tested. Given that both models did not show cytokinin responding to sucrose, these models should be considered questionable.

In order to ascertain differences between models that are valid for individual perturbations, it is necessary to observe the emergent properties of these networks as they relate to outcome landscapes (Figure 7B), and their specific prediction failures 7C). The patterns of mismatched biological and simulated outcomes for different networks can be used to judge features of the various models for biological validity. The two top performing models, Bertheloot and when a stimulatory edge has been added between sucrose and the branching outcome (Fig 7Ai and 7Aiii), were able to correctly predict the biological outcome when both sucrose and auxin were 0. Both these models failed once sucrose was increased slightly (*Aux* = 0, *Suc* = 0.2), suggesting that both models were unable to capture the immediate response that the plants had to any sucrose present in the system. Given the profile of the remaining mismatches, it is hard to state with any certainty which of the two models is better. To distinguish between these two models, experiments would need to be performed in the extreme high sucrose, high auxin portion of the simulated landscape (Fig 7Bi and iii).

When looking at the model that removed the stimulatory edge between sucrose and cytokinin (Fig 7Aii), this system predicts a low sensitivity to the presence of sucrose, which is counter to biological expectations, in addition to the already stated invalidity of failing to predict change in cytokinin in response to perturbing sucrose. The mismatches recorded consistently display that this model is unable to integrate the role of sucrose in producing the phenotypic outcome. An interesting set of models to compare are the ones in which a stimulatory edge has been added from sucrose to the branching outcome. Both models produced similar landscapes for the branching outcome by varying auxin and sucrose but have little overlap in which simulations failed to align with experiments. Of these, there are key areas that could be used to distinguish between the properties of each model. The first area of comparison is the behaviour of sucrose at extremely low values. The second area of comparison is the already stated response of cytokinin to sucrose.

## Discussion and Future Directions

The PSoup approach provides a versatile, simple to use, mathematically transparent approach to bridge the gap between mechanistic ODE type models and simple Boolean like models. Importantly, PSoup is designed to be testable against diverse data types and experimental conditions as demonstrated in this study. This means that data from current and historic sources can be integrated when testing the model. Meanwhile the flexibility of representation allowed means that a knowledge graph can be produced that incorporates interactions from multiple biological scales (including genes, proteins, broad phenotypes, and environmental interventions).

PSoup is useful as a tool for clarifying understanding, communicating concepts with others, and testing hypotheses. The construction of a diagrammatic representation of the system of interest provides an intuitive entry point for individuals to represent their understanding of the system, which can then be vetted without any need for the construction of mathematical equations, the establishment of kinetic parameters, or the need to make decisions on how information propagates through the system. This study has demonstrated that the PSoup method has a high degree of success in predicting the outcome of perturbations in a simple network model. For an example of the predictive power of PSoup in a complex network, see Mitsanis et al. (2026).

Once PSoup translates the diagram into a mathematical model, simulations can be run to test the logical outcomes of a system given the understanding of the user, and test conclusions against biological data. As such, we propose that PSoup is a tool that can be used by scientists (including students) to first clarify understanding and then assist in the creation of hypotheses to be tested. Of particular benefit is when there are multiple plausible ideas for the topology underlying a system. Separate diagrams can be produced depicting alternate topologies, which are then exposed to the same conditions for simulation. This can be used to identify conditions under which separate interpretations of the system are expected to behave differently.

Given the flexibility of the PSoup approach, this method is relatively easy to apply in entirely novel systems. Indeed, the method employed in the PSoup package is scalable to the level of understanding for a given system and can be updated readily as new information becomes available. The automatic nature of the translation from diagram to model makes PSoup accessible even to biologists who are uncomfortable with mathematics. Meanwhile, the transparent nature of the models produced means that they can be readily interrogated by mathematicians, and the models updated if desired. This provides simplicity and automation, with the full capacity to modify and refine to those who want it. This reduces the need to spend time, money, or computing power trying to establish ‘plausible’kinetic parameter values, which allows the user to focus on the ‘goal’of the network in question without overstating existing biological knowledge. Here, users can benefit from the ability to scale up the complexity of the network that they are exploring, while maintaining that each edge is grounded in knowledge taken from the literature (see Mitsanis et al. (2026)).

In addition to PSoup being beneficial for individual use, there is great potential to use it in conjunction with other existing models. The main goal of combining PSoup with other models is to incorporate an understanding of the underlying knowledge of the genetic and physiological network underpinning the system, potentially increasing the capacity to predict based on fundamental features of the system. The first connection being explored is to the Crop Growth Model APSIM (Keating et al., 2003;Holzworth et al., 2018). The goal of this connection is to explore cross scale modelling from gene network to consequences on crop yield. The second connection being explored is to Functional Structural Plant Models via the L-Systems formalism (Prusinkiewicz and Hanan, 2013). Such a connection would be useful to help explore the interaction between genotype and pruning/management practices on fruit yield. We suggest that there may be additional applications with Machine Learning, or possibly even AI, to guide these methods to more realistic answers by adding greater weight to connections that are known to be true. The ideal method for using PSoup in conjunction with AI is to design a Model Context Protocol (MCP) that can standardise how PSoup can be plugged into a variety of AI tools (Ray, 2025). This would facilitate the inclusion of PSoup in future endeavours while removing the need for new users to reconfigure the method for their own purposes.

For many of the proposed connections with other existing models, there is the challenge of making a meaningful connection between a model that is designed to be qualitative in nature, with a model that is quantitative. In these cases, it will be necessary to make modifications to the PSoup output by either establishing a heuristic to translate the output of PSoup to something that these quantitative models can understand, or there will need to be a reintroduction of parameters into PSoup that will allow the outputs to be applicable in the quantitative model. If the latter is pursued, the introduction of these parameters should be a last step once users are happy with the underlying topology of their system. Once that has been achieved, equations can be redesigned to have a form, including parameters, that achieve the correct ranking of outputs as seen in the biological data. This strategy preserves the philosophy of prioritising the importance of the network itself, with the fitting of the model existing only to improve the capacity to integrate with a quantitative model or prediction objective.

The main limitation of a PSoup model stems naturally from the lack of parameterisation. Without parameters to dictate the relative influence of different inputs on the outcome of a node, there will be issues when it comes to establishing when a system will transition between different categorical types, particularly when more than one conflicting perturbation is enacted simultaneously. In terms of reproducing qualitative observations, however, Lawson et al. (2026) found that a qualitative model and its fully-parameterised equivalent (calibrated to try to match those observations) were both similarly accurate at making predictions in general, with each performing better in different areas. While different algorithmic rules can and should be explored, without parameters this limitation will be constitutive. Within the scope of this study, as the network was simple and the number of conditions for simulation limited, we did not need to consider the use of a threshold for determining when a simulation implies a categorical change. However, as the size of the network and number of comparisons increases, a threshold above and below 1 may need to be selected as a criteria for when a category change from the baseline has been achieved. This threshold can even be optimised from the data used to validate the model. See Mitsanis et al. (2026) for an example.

It is important to recognise that despite the difficulty in establishing thresholds of categorical change, in particular when opposing interventions have been applied, PSoup still manages to maintain the trends apparent in developmental and physiological biology. If double perturbations are compared with their individual perturbation counterparts, PSoup is generally able to identify intermediate, or additive traits, excluding the existence of masking or epistatic effects (Supplementary Three). In fact, incorrect predictions for double perturbations can themselves be useful for identifying the relative importance of different inputs to the phenotypic outcome. Given the relative-to-baseline approach for defining simulations, it is natural to make this kind of conclusion. For example, if an opposing set of perturbations has been made, each modified by the same amount (eg. auxin modifier set to 2, and sucrose modifier set to 2), if one perturbation overwhelms the effect of another, it can be said the corresponding node has a greater influence on the system than the other. This information could be useful to produce more refined models in the future.

An additional difficulty that can arise from using models built with PSoup involves the decisions that need to be made regarding the values applied to perturbations. The first complication in this regard relates to the choice of baseline. Given that the values provided to modifiers that define perturbations are in relation to the baseline values, as well as the the outcomes of simulations, baseline choice itself is important. Choosing the wrong baseline could obfuscate incorrect model behaviour, or potentially even lead to a greater number of errors being reported than exist if the chosen baseline behaved unusually in the original experiment. Once the baseline has been established, this will be used to define the relative modifier values for all simulated perturbation types. However, it should be acknowledged that there may be questions regarding the ability of the plant to even absorb the totality of the intervention applied. Therefore, modifier values should be chosen carefully, and considered to be a best approximation given the knowledge available.

The formulation of PSoup as communicated here is simple. However, there are several extensions not explored in this paper that can allow for greater biological realism without betraying the design principle behind PSoup. The first extension relates to the breadth of the algorithmic rules themselves. The ability to explicitly provide an exogenous supply of some substance to a node (as opposed to just adjusting the modifier) will allow for users to perturb the system in multiple ways that may more accurately represent certain experimental treatments. Another extension that is included with PSoup but was not explored within this paper is the ability to choose the functional form that is applied to modifiers or necessary stimulants (for a discussion on possible functional forms and their biological interpretation see Supplementary Five). The base form that PSoup employs for both is linear. However, this may not be the best choice for representing biological interactions. For example, Michaelis–Menten kinetics are often employed to represent biochemical reactions (Cornish-Bowden, 2015).

Some extensions that could assist users in exploring the network itself would allow for the automatic generation of alternate networks that can be validated and compared with the same set of defined simulations. In the simplest case, an edge of type ‘unknown’(which exists in SBGN-AF language) could be used if there is an expected connection between two nodes. PSoup could then generate alternative networks, representing each possible edge type (and direction of causality) connecting those two nodes. Another possibility would be to take a starter network as an input and to generate each neighbouring network so that they can be compared. Such an extension could be the beginning of an optimisation algorithm for finding the best network to explain biological outcomes or to determine whether alternative networks could also explain the biological data. However, this style of use would be counter to the original intention of PSoup which is to explore bespoke network diagrams based on prior knowledge.

The PSoup approach discussed in this paper is both simple and powerful, with many opportunities to be adapted and extended. Given that much of scientific activity is increasing driven by large amounts of data and heavy computing, PSoup represents an opportunity to simplify, and make the most of existing knowledge of system structure in our attempts to solidify understanding.

## Supporting information

Supplementary One

Supplementary Two

Supplementary Three

## Abbreviations

ODE: Ordinary Differential Equation
SBGN: Systems Biology Graphical Notation
AF: Activity Flow

## Author contributions statement

N.Z.F, C.A.B, B.A.J.L and K.B were all involved in the initial conception of this project. N.Z.F designed and built the R package with substantial insights from B.A.J.L. and C.M. N.Z.F ran the simulations and analysed the results. N.Z.F wrote the majority of the text with contributions from C.A.B. B.A.J.L contributed text and simulations associated with the Bayesian analysis of the original Bertheloot model, and wrote the supplementary materials comparing the PSoup approach to ODEs, and discussing the functional forms of modifiers and necessary stimulants. C.A.B, B.A.J.L, C.M and K.B provided feedback on the text.

## Acknowledgments

The authors thank Dr. Jim Hanan for extensive feedback he provided on the text. We also thank Dr. Nicholas O’Brain for feedback provided in the final drafting of this work. This work is supported by funds from the ARC Centre of Excellence for Plant Success in Nature and Agriculture,.

