## Supplementary Materials for "PSoup: an R package for simulating biological networks from a qualitative perspective"

### Supplementary One

An Excel spreadsheet summarising experiments from Bertheloot (Sup1.xlsx) is included.

The following results from the Bertheloot paper were excluded from the summary to prevent duplications or remove data for which a comparison was not tractable. Figure 2d was excluded as it represented the same outcomes as found in Figure 2c, simply recording a slightly different measure that tracked the same underlying branching phenotype with no deviation in outcome. Figure 2f (not in excel sheet) was excluded as the data was too messy to interpret. For Figure 3a, the intermediate forms of CK (not in excel sheet) were excluded from the analysis as they followed the same trend as the active form of CK and had no separate node to track in the PSoup diagram and therefore represent repeats. Figures 5b and 5c were also excluded as they represented composite figures that took datapoints from other figures. These exclusions reduced the originally considered experimental conditions from 122 to 78.

**Table 1** A summary of the conditions considered to represent the natural baseline for each node in the PSoup Bertheloot model for the Rose and Pea systems.

| Node | Rose baseline | Pea baseline |
|---|---|---|
| Auxin | Intact shoot/1mM NAA | Intact shoot/1mM NAA |
| Sucrose | Intact leaves/50mM Sucrose | Intact leaves/70mM Sucrose |
| Sucrose | WT/-SL | WT/-SL |
| Cytokinin | WT/-CK | WT/-CK |

The natural baseline often did not exist in each figure, and so a surrogate baseline was chosen. Highlighting was used to indicate which conditions within each figure was chosen to represent the baseline. Highlighting was also used to indicate whether non-baseline conditions represented an increase or decrease from the baseline, whether the values recorded represent the recorded phenotypic outcome of the experiment, and the biological system of the experiment. The spreadsheet contains a key for this information.

### Supplementary Two

An Excel spreadsheet showing the translation of the data found in Supplementary One into the definition file for running simulations (Sup2.xlsx).

The rows of data in this spreadsheet correspond to the conditions found in Supplementary One. The values are simply normalised against the selected baseline for each figure. The columns of this supplement (excluding the Fig column) relate to the modifiers of the model and therefore define the perturbations to be simulated. The highlighting indicates the baseline within each figure set, which by definition will have values only of 1. The information in this file was used to fill the modifierDef file generated by PSoup.

### Supplementary Three

An Excel spreadsheet showing the final values generated by simulations run based on the information found in Supplementary Two (Sup3.xlsx).

The rows for data in this spreadsheet correspond to the conditions found in Supplementary One and Supplementary Two.

### Supplementary Four

This supplement gives a demonstration on the comparative behaviour of experimental and simulated data seen in Figure 5 of the main text, the experimental data from the Bertheloot 2019 paper is compared with the simulated outcome produced using the PSoup method. In that comparison, each data point was compared with a selected baseline chosen within each figure of origin. The comparison showed only if the results were equal to, above, or below that of the baseline condition. While this categorical comparison is appropriate given that the PSoup method is free of kinetic parameters, it limits a full understanding of what is going on both in the biology and the simulations, particularly in the case that multiple perturbations are occurring at once.

When making a physiological comparison of some phenotype in a set of plant experiments, it is rare that all outcomes are compared only to the control condition. Multiple comparisons are needed, especially to reveal epistatic effects. In the case of a gradient of interventions (perturbations), each intervention will be compared with its neighbours to see if the effect on the phenotype is also a gradient, or if the effect plateaus. In the case that multiple interventions have been applied on a single plant, that plant will be compared with plants that exhibit each individual intervention. In such a comparison, multiple types of dynamics can be revealed. In the case of antagonistic perturbations, it can be seen if the resultant phenotype is intermediate to the individual perturbations, or if the system becomes dominated by one of the perturbations. In the case of multiple perturbations of concordant effect, it can be revealed if combining the perturbations results in an additive effect on the phenotype, or if the phenotype is dominated by one of the perturbations.

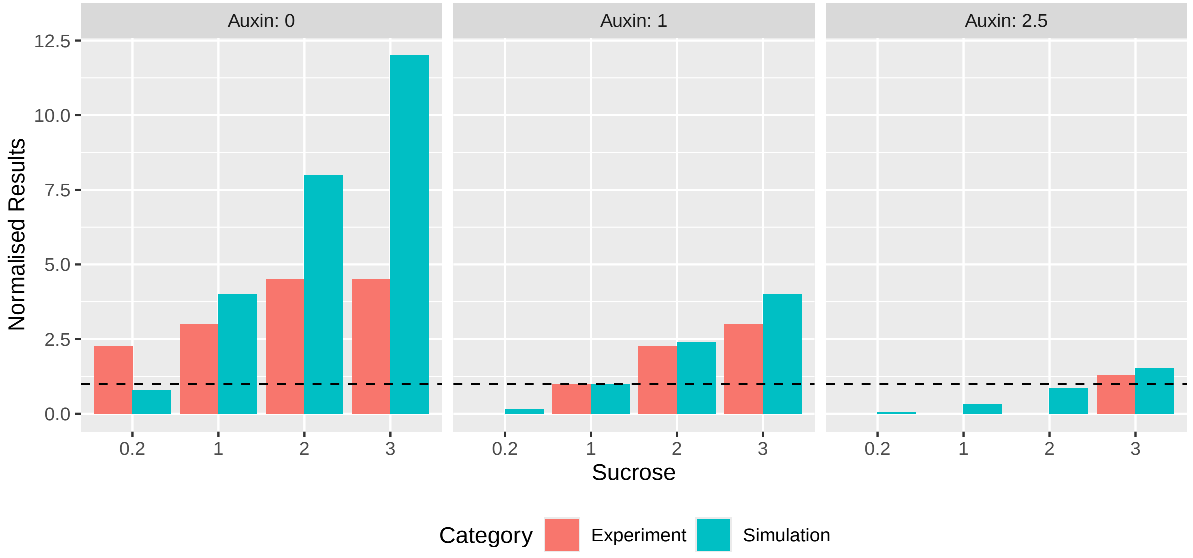


**Figure 1** Supplementary Figure : A comparison of the biological results of figure 2c of the Bertheloot paper, and their PSoup recreations. Panels are organised by level of Auxin, with each demonstrating the effect of Sucrose at that Auxin level. Orange bars represent the normalised experimental results, teal represents the PSoup outcome. Values of Auxin and Sucrose and Auxin are expressed relative to the selected baseline of the figure. The baseline values are indicated with a horizonal dashed line for ease of comparison. Values above this line correspond to the increase category of Figure 4, while values below this value correspond to the decrease category.

Given that the PSoup method does not use kinetic parameters, the results returned are qualitative in nature. This means that the magnitude of change indicated by PSoup is not reliable and instead should only be used to predict the direction of change. This also means that PSoup will have trouble predicting the boundaries (particularly in the case of multiple perturbations) for when the system transitions from one category to another. Even so, in the case of single perturbations considered at a time, PSoup can be relied upon to indicate the correct direction of change given an accurate diagrammatic representation of the system.

Supplementary Figure 1 shows a subsection of the results seen in Figure 5 of the main text, in particular that relating to Figure 2c from Bertheloot et. al., (2019). Below we compare the normalised experimental results, with that of their simulated PSoup counterparts. In each case, the selected baseline condition is represented by 1. In this figure it can be seen that within each auxin grouping that the trend in both experiments and simulations is an increase in the branching phenotype as the amount of sucrose in the system increases. In addition, if sucrose conditions are compared between the auxin panels, the trend for both experiments and simulations is that as auxin increases, the normalised branching decreases. This behaviour of PSoup matching the direction of trend in one direction is generally consistent across all data analysed in this paper, apart from some questionable biological results already discussed in the main text. An example of one of these is included in Figure 1, when Aux = 0 and Suc = 0.2 (the first set of bars). This mismatch between the biology and the simulation relates to their position above or below the baseline value 1 (indicated with a dashed line). The biology indicates an increase from the baseline, while the simulation indicates a decrease. Even so, the trend between the experiments and the simulations is maintained.

#### *Example Figure demonstrating trend*

Figure 2c from the Bertheloot was chosen as an example, not only because it demonstrates an organised exploration of two interventions, but also because it reveals certain quirks of the comparison between experiments and simulations. First, is the stated unreliability regarding the prediction of magnitude. One consequence of this is that PSoup will struggle to capture natural thresholds in the biological data. This can be seen in the ‘Auxin: 0’ panel where the biology shows no further increase in normalised branching outcome beyond a sucrose level of 2. Another example of this thresholding is in the ‘Auxin: 1’ and ‘Auxin: 2’ panels, for the lower sucrose values. In these experiments, the branching phenotype never reached the length at which it could be measured and so the raw data recorded an NA value. The conclusion that these conditions have a reduced branching phenotype is still valid, PSoup simply does not reflect the same graded decrease in phenotypic outcome.

#### *Remaining data used in paper comparing normalised biological values with simulated outcomes*

The following figures (in addition to Figure 1 above), show the normalised biological results, and raw simulated outcomes seen in Figure 5 of the main text. In cases where the experimental outcomes and simulation are not aligned, some commentary will be given. Otherwise, only the figure is presented.

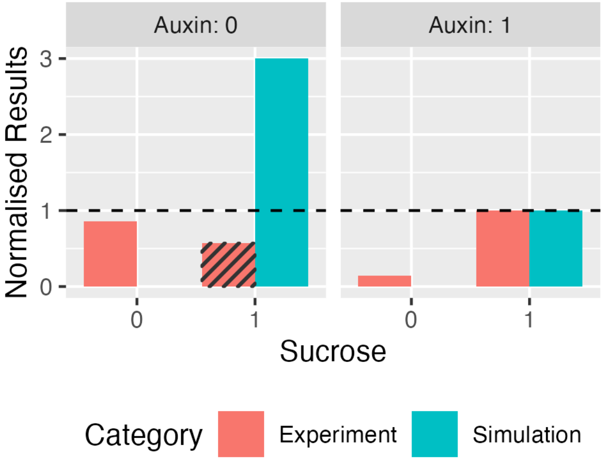


**Figure 2** Data from the Figure 2a subsection of Figure 5 of the main text.

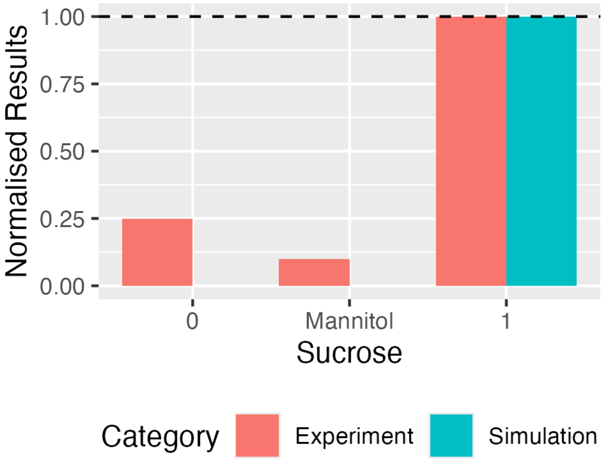


**Figure 3** Data from the Figure 2b subsection of Figure 5 of the main text.

Supplementary Figure 2 corresponds to Figure 2a within Figure 5 of the main text. This figure shows an example of a mismatch between the experimental and simulated data. As explained in the main text, the experimental condition indicated with cross hatching is an example of a biological outcome that is not behaving as expected. Not only is this outcome inconsistent with other experimental outcomes for this tested condition, but this particular result was also not statistically significantly different from the baseline condition.

Supplementary Figure 6, which corresponds to the information in subsection Figure 3b of Figure 5 in the main, recapitulates the mismatch where the biology did not increase with increasing sucrose as predicted in the simulation.

Supplementary Figure 9 corresponds to the Figure 4b subsection of Figure 5 in the main text. Here we see the remaining two mismatches between the experiments and simulations, specifically when the biological result is in question (cross hatching). Both these matches are one of a set representing the same condition. They represent the only instances of these conditions not behaving as predicted by the simulations.

Supplementary Figure 11 represents the Figure 4d_a subsection of Figure 5 of the main text. It demonstrates the mismatch in the main text where the simulation expected an increase relative to the baseline when strigolactone was unavailable in the system. Instead the experimental outcome matched that of the baseline.

Supplementary Figure 13 represents the Figure 4e subsection of Figure 5 of the main text. It depicts the mismatch where the simulation predicted an increase relative to the baseline, when the biology showed a decrease. Biologically speaking, the system stayed the same under the ‘SL = 0, and CK = 0.5’ scenarios even when sucrose was added to the system.

Supplementary Figure 14 represents the Figure 4f subsection of Figure 5 of the main text. It depicts the mismatch where the simulation predicted an increase relative to the baseline, when the biology showed a decrease. Biologically speaking, the

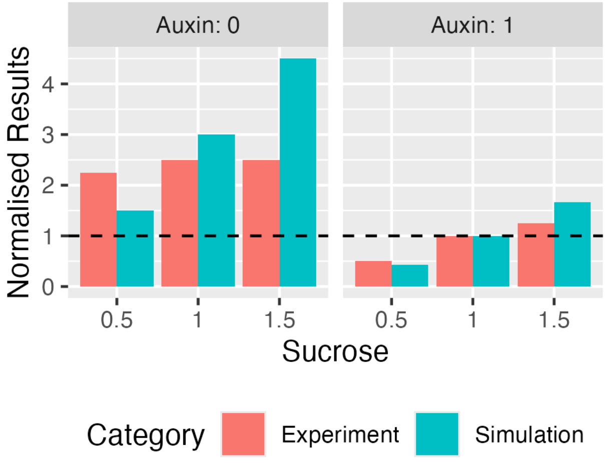


**Figure 4** Data from the Figure 2e subsection of Figure 5 of the main text.

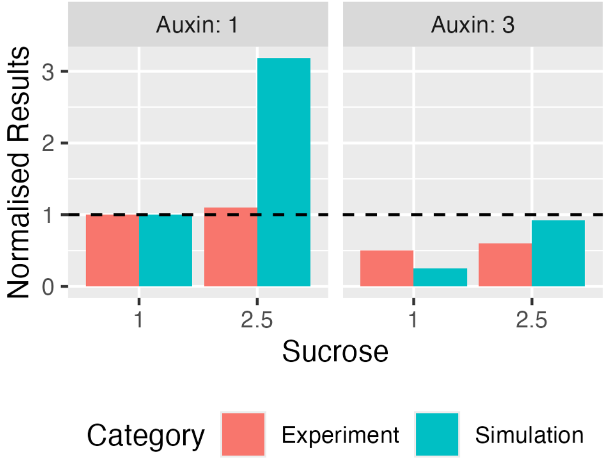


**Figure 5** Data from the Figure 3a subsection of Figure 5 of the main text.

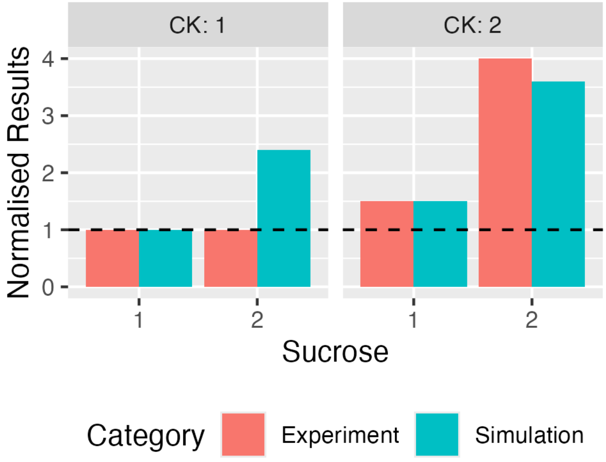


**Figure 6** Data from the Figure 3b subsection of Figure 5 of the main text.

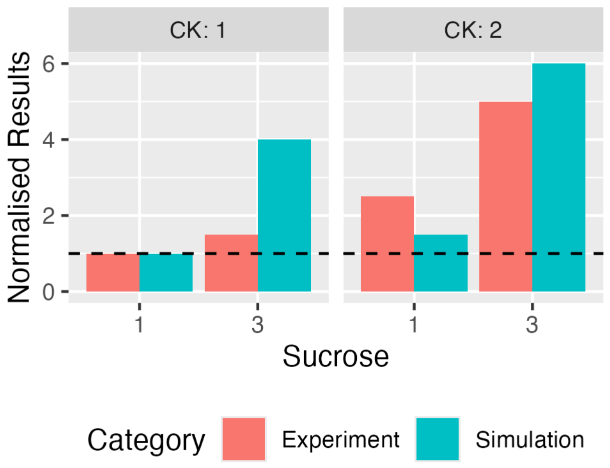


**Figure 7** Data from the Figure 3c subsection of Figure 5 of the main text.

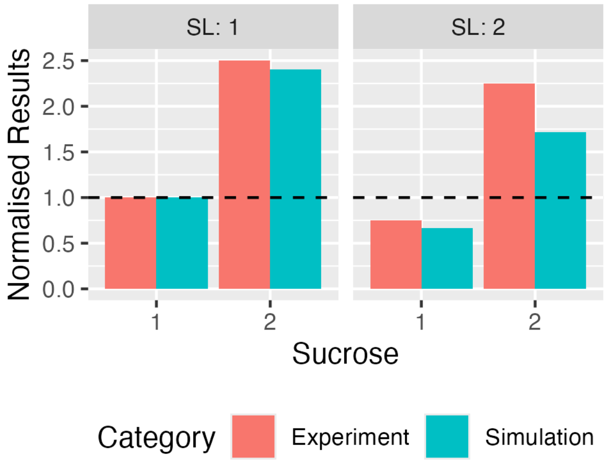


**Figure 8** Data from the Figure 4a subsection of Figure 5 of the main text.

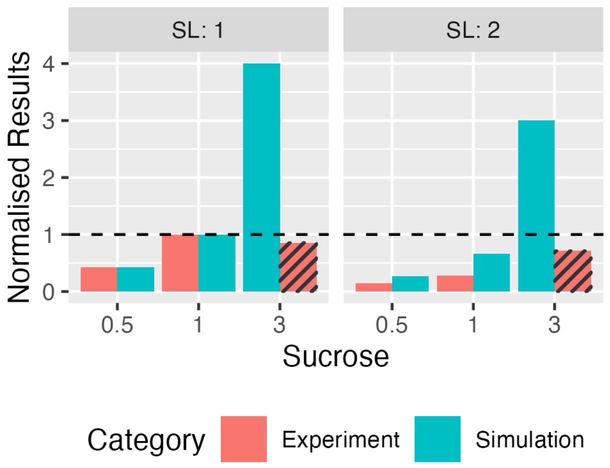


**Figure 9** Data from the Figure 4b subsection of Figure 5 of the main text.

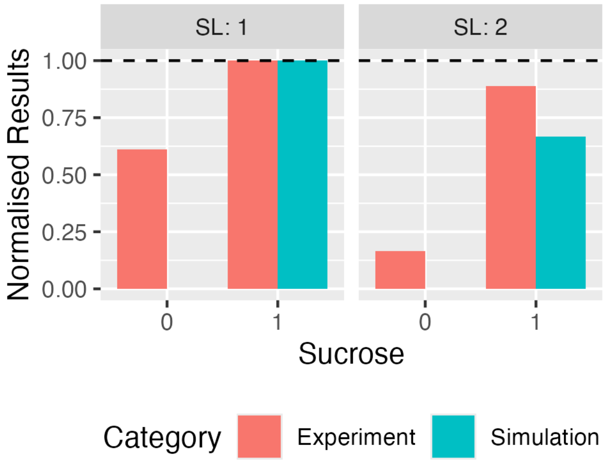


**Figure 10** Data from the Figure 4c subsection of Figure 5 of the main text.

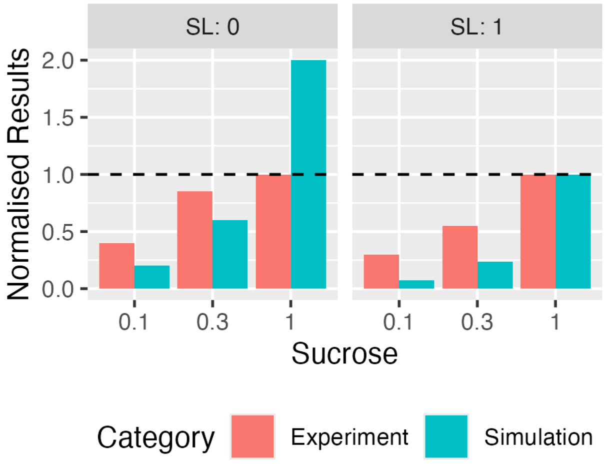


**Figure 11** Data from the Figure 4d_a subsection of Figure 5 of the main text.

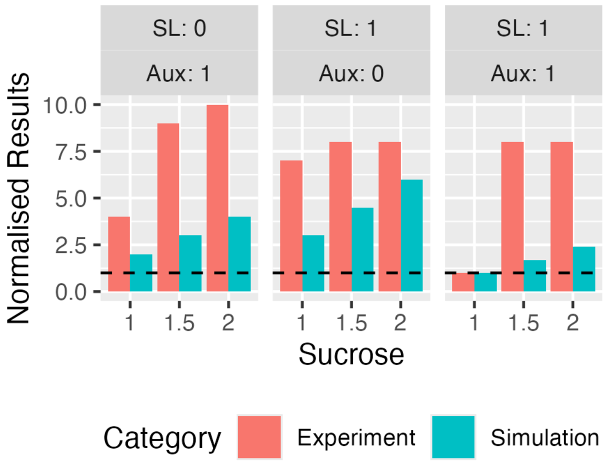


**Figure 12** Data from the Figure 4d_b subsection of Figure 5 of the main text.

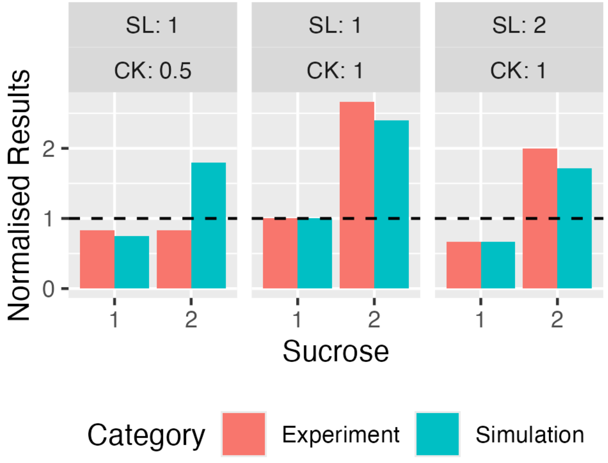


**Figure 13** Data from the Figure 4e subsection of Figure 5 of the main text.

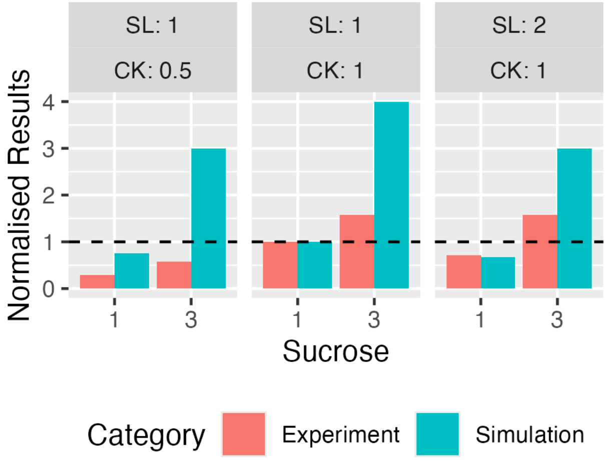


**Figure 14** Data from the Figure 4f subsection of Figure 5 of the main text.

system under the 'SL = 0, and CK = 0.5' scenario only had a minor increase in branching when sucrose was added to the system.

#### *Supplementary Four methods*

To produce Supplementary Figure 1, the raw experimental data for Figure 2c of the Bertheloot paper was collected along with the corresponding simulated data. The simulated data was already appropriate for plotting, however the experimental data needed to be processed to allow for comparison. The first step for processing required that the value for the selected baseline condition needed to be used to normalise all the values from this experiment so that outcomes were expressed relative to the baseline 1.

The biological data for this figure measured the time taken for the bud to reach a length of 4mm. This means that the smaller the value recorded, the higher the branching value. Therefore, the direction of the values relative to one needed to be reversed to be compared with the simulated data. First the log was taken for all the branching outcomes. This made all results relative to 0. Next these values were multiplied by -1 to reverse the direction of the results relative to 0. Finally, the exponential was taken to reestablish the baseline relative to 1.

Supplementary Figure 2 to Supplementary Figure 14 were easier to produce as they did not need a reversal of the direction indicated by the normalised biological results. In all other respects, the data of these figures was treated the same as that of Supplementary Figure 1.

### Supplementary Five

The Bertheloot model describes the dynamics of cytokinin ($CK$) and strigolactone ($SL$), as well as their effect on branching inhibition through an integrated signalling factor ($I$), as a series of ordinary differential equations,

$$\frac{dCK}{dt} = \frac{c_1}{1+b_1 A} + \frac{a_1 S^2}{k_1 + S^2} - d_1 CK$$
$$\frac{dSL}{dt} = c_2 + \frac{a_2 A^2}{k_2 + A^2} - d_2 SL$$
$$\frac{dI}{dt} = c_3 + \frac{a_3 SL^2}{1+(\mu_1 + \mu_2 S^2)SL^2} + \frac{a_4}{1+k_3 CK^2} - d_3 I$$

Lowercase variables are the parameters, and capture biochemical kinetic constants. The dynamics are also modulated by the levels of auxin ($A$) and sucrose ($S$), which are treated in the model as constant.

Matching the 2019 Bertheloot study, the data are treated as corresponding to the equilibrium reached by the system, which is given by setting the three rates of change to zero,

$$CK^* = \frac{c_1}{d_1}\frac{1}{1+b_1 A} + \frac{a_1}{d_1}\frac{S^2}{k_1 + S^2}$$
$$SL^* = \frac{c_2}{d_2} + \frac{a_2}{d_2}\frac{A^2}{k_2 + A^2}$$
$$I^* = \frac{c_3}{d_3} + \frac{a_3}{d_3}\frac{SL^{*^2}}{1+(u_1 + u_2 S^2)SL^{*^2}} + \frac{a_4}{d_3}\frac{1}{1+k_3 CK^{*^2}}$$

This motivates choosing new parameters,

$$\lambda_{cka} = \frac{c_1}{d_1}, \quad \lambda_{cks} = \frac{a_1}{d_1}, \quad \lambda_{sl} = \frac{c_2}{d_2}, \quad \lambda_{sla} = \frac{a_2}{d_2}, \quad \lambda_i = \frac{c_3}{d_3}, \quad \lambda_{isl} = \frac{a_3}{u_1 d_3}, \quad \lambda_{ick} = \frac{a_4}{d_3}$$

$$K_{cka} = \frac{1}{b_1}, \quad K_{cks} = \sqrt{K_1}, \quad K_{sla} = \sqrt{K_2}, \quad K_{is} = \sqrt{\frac{u_1}{u_2}}, \quad K_{isl} = \sqrt{\frac{1}{u_1}}, \quad K_{ick} = \sqrt{\frac{1}{K_3}}$$

that allow the equilibrium values for the state variables to be written

$$CK^* = \lambda_{cka}\frac{K_{cka}}{K_{cka} + A} + \lambda_{cks}\frac{S^2}{K_{cks} + S^2}$$
$$SL^* = \lambda_{sl} + \lambda_{sla}\frac{A^2}{K_{sla}^2 + A^2}$$
$$I^* = \lambda_i + \lambda_{isl}\, f(S)\,\frac{SL^{*2}}{K_{isl}^2\, f(S) + SL^{*^2}} + \lambda_{ick}\frac{K_{ick}^2}{K_{ick}^2 + CK^{*^2}}$$
$$f(S) = \frac{K_{is}^2}{K_{is}^2 + S^2}$$

This now expresses the steady state of the system in terms of the maximal possible contributions of each term, $\lambda$, and half-saturation constants, $K$. It is the values of these thirteen constants for which Bayesian inference is performed.

Reflecting a lack of quantitative information regarding the values of the model's parameters, we assign them a non-informative prior distribution that allows them to potentially vary across a broad range of values. This prior is uniform in scaling (log-uniform in value) such that doubling or halving their value is assigned the same weight,

$$\log_{10}\lambda \sim U[-5,5], \quad \log_{10} K \sim U[-5,5]$$

The original data are given in the form of measurements ($CK^*, SL^*, T$), corresponding to different auxin/sucrose conditions, $\mathbf{x} = (A,\ S)$. Here $T$ denotes the time until significant branch elongation, but the model predicts an equilibrium level of anti-budding factor, $I^*$. We convert the observed budding times into (hypothetically measured) levels of anti-budding factor using the relationship given in the 2019 Bertheloot study,

$$I^* = \frac{T+2.2}{3.5} \qquad \text{if } T < 8.3$$
$$I^* \geq 3 \qquad \text{if } T \geq 8.3$$

This gives us data $\mathbf{y} = (CK^*, SL^*, I^*)$ that match the nature of the model's predictions. We link observations and model predictions using a standard statistical model with Gaussian errors (Lambert et al., 2023),

$$\mathbf{y} = \mathbf{M}(\mathbf{x};\boldsymbol{\theta}) + \boldsymbol{\epsilon}, \quad \epsilon_i \sim \mathcal{N}(0, \sigma_i^2)$$

where $\mathbf{M}(\mathbf{x};\boldsymbol{\theta})$ denotes evaluating the model with experimental conditions $\mathbf{x}$ and values $\boldsymbol{\theta}$ for the thirteen parameters, and $\mathcal{N}(\mu, \sigma^2)$ denotes the univariate normal distribution with mean $\mu$ and variance $\sigma^2$. Given the small amount of data available for calibration, we do not try to also learn the standard deviations of the errors associated with taking measurements of cytokinin ($\sigma_{ck}$), strigolactone ($\sigma_{sl}$) and inhibitory factor ($\sigma_i$). Instead, we fix these at values calculated as 10% of the standard deviation of the observed data for each species, leaving plenty of variation for the model and calibration process to attempt to explain. The choice not to infer the level of measurement noise also aids the inference process, by avoiding trapping it in a state where the posterior tends to favour explaining a lot of the variance in the data using measurement noise.

Given that the chosen error model treats the errors on the three model variables as independent, the likelihood for one experiment is given by their product. However, we must also take into account that any amount of inhibitory factor at a level $I = y_3 > 3$ will be recorded as a failure to bud. Thus, for experiments where budding failure was the recorded result, any values $y_3 = I^* + \epsilon_3 > 3$ need to be treated as agreeing with the data. In statistics, this is known as censored data, and the likelihood for these measurements is calculated by integrating the probability density function over the region that exceeds the threshold. This results in an overall likelihood for each individual experiment that takes the standard form

$$p(\mathbf{y}|\mathbf{x},\boldsymbol{\theta}) = \frac{1}{\sigma_{ck}\,\sigma_{sl}\,\sigma_i}\phi\left(\frac{y_1 - M_1(\mathbf{x};\boldsymbol{\theta})}{\sigma_{ck}}\right)\phi\left(\frac{y_2 - M_2(\mathbf{x};\boldsymbol{\theta})}{\sigma_{sl}}\right)\phi\left(\frac{y_3 - M_3(\mathbf{x};\boldsymbol{\theta})}{\sigma_i}\right)$$

if the experiment had a budding time recorded, and

$$p(\mathbf{y}|\mathbf{x},\boldsymbol{\theta}) = \frac{1}{\sigma_{ck}\,\sigma_{sl}}\phi\left(\frac{y_1 - M_1(\mathbf{x};\boldsymbol{\theta})}{\sigma_{ck}}\right)\phi\left(\frac{y_2 - M_2(\mathbf{x};\boldsymbol{\theta})}{\sigma_{sl}}\right)\left(1 - \Phi\left(\frac{3 - M_3(\mathbf{x};\boldsymbol{\theta})}{\sigma_i}\right)\right)$$

if no budding was recorded. Here $\phi(z)$ denotes evaluation of the unit normal density function at point $z$, and $\Phi(z)$ denotes evaluation of the unit normal cumulative density function. We treat the results of the $N$ experiments as uncorrelated with one another, such that the likelihood for the whole dataset $\mathbf{d}$, consisting of experiments with conditions $\mathbf{x}_i$ and observations $\mathbf{y}_i$, is the product of the likelihoods given above,

$$p(\mathbf{d}|\boldsymbol{\theta}) = \prod_{i=1}^{N} p(\mathbf{y}_i|\mathbf{x}_i,\boldsymbol{\theta})$$

Combining the prior and likelihood using Bayes' theorem gives us the non-normalised posterior density

$$p\left(\boldsymbol{\theta}|\mathbf{d}\right) \propto p(\boldsymbol{\theta})p\left(\mathbf{d}|\boldsymbol{\theta}\right),$$

that we generate samples from using sequential Monte Carlo (SMC) (Gilks and Berzuini, 2001; Neal, 2001; Drovandi and Pettitt, 2011). SMC generates and manipulates a specified number of particles (here $N_{\text{particles}} = 20,000$) that represent samples from a sequence of distributions that begins at the prior and transitions geometrically to become the posterior distribution,

$$p_\gamma(\boldsymbol{\theta}|\mathbf{d}) \propto p(\boldsymbol{\theta})[p(\mathbf{d}|\boldsymbol{\theta})]^\gamma \qquad \gamma \in [0,1].$$

The algorithm begins by taking $N_{\text{particles}}$ samples from the prior, generating particles corresponding to $\gamma_1 = 0$. Then, an updated value for $\gamma_2$ is decided by selecting how much of the complexity of sampling the actual posterior (which corresponds to $\gamma = 1$) should be introduced, given the quality of the current particles in terms of being samples from an updated distribution with $\gamma_{n+1} = \gamma_n + \Delta\gamma$. This judgement is made using the (un-normalised) importance weights of the particles with respect to a proposed new distribution relative to the current one, which are given by

$$w_i(\Delta\gamma) = \frac{p(\boldsymbol{\theta})\,[p(\mathbf{d}|\boldsymbol{\theta})]^{\gamma_n+\Delta\gamma}}{p(\boldsymbol{\theta})\,[p(\mathbf{d}|\boldsymbol{\theta})]^{\gamma_n}} = [p(\mathbf{d}|\boldsymbol{\theta})]^{\Delta\gamma}.$$

The quality of particles is evaluated using the effective sample size ($ESS$),

$$ESS(\Delta\gamma) = \frac{\left[\sum_{i=1}^{N_{\text{particles}}} w_i(\Delta\gamma)\right]^2}{\sum_{i=1}^{N_{\text{particles}}} w_i^2(\Delta\gamma)},$$

with the update at each iteration $\Delta\gamma$ decided by selecting the maximum value that does not exceed one (the posterior) and does not cause $ESS(\Delta\gamma)$ to fall below some fraction of the total number of particles, $\alpha N_{\text{particles}}$. The algorithm parameter

$0 < \alpha < 1$specifies how conservative the algorithm is, with larger values of $\alpha$ forcing the effective sample size to remain larger and hence increments $\Delta\gamma$ to be smaller. Here, $\alpha = 0.75$. Once a value of $\Delta\gamma$ has been decided, the particles are resampled using multinomial resampling according to their weights. This causes the particles to all become equally weighted again, at a cost of some duplication of higher-weight particles and some elimination of lower-weight particles. To then correct for duplication, several rounds of standard Markov chain Monte Carlo (MCMC) are applied to the entire particle set, targeting the new distribution, $p_{\gamma+\Delta\gamma}(\boldsymbol{\theta}|\mathbf{d})$. These MCMC steps are performed using multivariate normal random jumps, where the covariance of the $\boldsymbol{\theta}$ values across the particle set, $\Sigma$, is used to inform the scale of the jumping distribution,

$$\boldsymbol{\theta}'|\boldsymbol{\theta} \sim N\left(\boldsymbol{\theta}, \frac{2.38^2}{d}\Sigma\right),$$

where $d = 13$ denotes the dimension of the parameter space. The number of MCMC steps to perform at each iteration $n$ is chosen adaptively by first estimating their rate of acceptance, $A$, using five pilot MCMC steps and then enacting up to twenty-five more according to the formula used by Drovandi and Pettitt (2011),

$$R_{\text{extra}} = \min\left(\hat{R}, 25\right), \qquad \hat{R} = \max\left(0, \frac{\ln 0.05}{\ln(1-A)} - 5\right).$$

Note that $A$is first clamped to the range $\left[10^{-6}, 1 - 10^{-6}\right]$ to avoid the cases $A = 0$, that results in division by zero, and $A = 1$, which results in taking the logarithm of zero.

SMC repeats the process laid out above, iteratively choosing a new value for $\gamma_n$ by considering how much of an increment $\Delta\gamma$ is statistically "safe" (in terms of maintaining sufficient $ESS$), resampling to get unweighted particles from the new distribution, and then jittering these particles using MCMC steps to regain particle uniqueness. This repeats until the value of $\gamma = 1$ is reached, at which point the final iteration's resampling and jittering steps produce a set of $N_{\text{particles}}$ unweighted samples from the posterior distribution, the majority of which should be unique. These samples represent different possible calibrations of the model, which are used in the main manuscript to demonstrate the uncertainty that remains in the system's behaviour even after quantitative calibration to data.

### Supplementary Six

PSoup is a qualitative modelling approach, but is directly inspired by quantitative descriptions of the chemistry that governs biological networks. Here, we demonstrate how the functional forms chosen to define the different types of network edge in a PSoup model can be derived from biochemical arguments under certain assumptions. We begin with a standard differential equation approach for describing biochemical kinetics, where the concentration of each species varies with time according to

$$\frac{d[C_i]}{dt} = \lambda_i f_i([\mathbf{C}]) - \delta_i [C_i]$$

where $[C_i]$ denotes the concentration of the $i$-th chemical species, $\lambda_i$ denotes the overall production rate of this species, $\delta_i$ denotes the degradation rate of the species, and the function $f_i([\mathbf{C}])$ describes how the concentrations of all species (gathered together with some slight abuse of notation as vector $[\mathbf{C}]$) influence the production rate. As the definition of $f_i$ is left general, the only assumption made to this point is that all species undergo first-order degradation at some rate.

Our interest in a PSoup model is in how the system of interest behaves relative to some notion of baseline, representing for example the wild type. As the important behaviour of biological networks is often their steady state (Dun et al., 2009; Schaffter et al., 2011), including here, we choose the steady state of the system of equations of the form above as the baseline values. Define the functions $f_i([\mathbf{C}])$ such that they take on a value of unity at the baseline equilibrium state, which can always be done by re-scaling them and then absorbing the effect of this scaling into the definition of $\lambda_i$. Then, representing equilibrium by setting rates of change to zero, we have that for the concentration of each species, $[C_i]$, its baseline equilibrium value is $s_i = \lambda_i/\delta_i$.

Now we introduce non-dimensional variables that represent the levels of the network species relative to baseline, $C_i = [C_i]/s_i$, and associated vector of all levels, $\mathbf{C} = (C_1, C_2, \ldots)$. Substituting into the differential equation above,

$$\frac{dC_i}{dt} = \delta_i\big(f_i^*(\mathbf{C}) - C_i\big),$$

where $f_i^*(\mathbf{C})$ denotes evaluating the function $f_i([\mathbf{C}])$ after first scaling each element of the input by its baseline value. That is, $f_i^*(C_1, C_2, C_3, \ldots) = f_i(s_1C_1, s_2C_2, s_3C_3, \ldots)$. Note that the non-dimensionalised variables $C_i$ are those used in the PSoup description of a network, taking on a value of one under baseline/wild-type conditions and varying under perturbed scenarios relative to this. Similarly, the condition on the function $f_i$ that it takes on a value of unity at baseline simply becomes $f_i^*(\mathbf{1}) = 1$ for its equivalent in the non-dimensionalised description, where $\mathbf{1}$ denotes the vector of all ones.

To create a simplified and qualitative model, PSoup assumes that the inherent timescales for the dynamics of all of the non-dimensionalised chemical species are equal, $\delta_i = 1/\tau$ for all $i$. Then, by approximating the dynamics of this system of differential equations using a single forwards Euler step over timeframe $\tau$,

$$\begin{aligned} C_i(t+\tau) &\approx C_i(t) + \tau \cdot \frac{1}{\tau}\left(f_i^*\big(\boldsymbol{C}(t)\big) - C_i(t)\right) \\ &\approx f_i^*\big(\boldsymbol{C}(t)\big) \end{aligned}$$

The updates used by PSoup fit into this form, and thus concord with biochemical dynamics for which the degradation rate of all species is treated as equal, but relative synthesis rates and hence actual concentrations at equilibrium are unconstrained. For the remainder of the discussion here, we drop the star notation for simplicity.

To ensure the property $f_i^*(\mathbf{1}) = 1$ is satisfied and hence provide a PSoup model with the specific property of being defined relative to a baseline, the synthesis rate modifier functions are constructed using a product of separate functions that all themselves satisfy $f(\mathbf{1}) = 1$,

$$f_i^*(\boldsymbol{C}) = f_{i,\text{stim}}(\boldsymbol{C}) \cdot f_{i,\text{inhib}}(\boldsymbol{C}) \cdot f_{i,\text{necstim}}(\boldsymbol{C}).$$

Taking the product of functions in this manner can be interpreted as treating the effects of stimulation, inhibition and necessary stimulation independently. To illustrate this using an example, let us denote by $X$ some single species in a network, the synthesis of which is stimulated by a single species $S$, and inhibited by a single species $I$. Then, the update formula generated by PSoup (as laid out in the main document, and justified below) is

$$X(t+\tau) = S \cdot \frac{2}{1+I}$$

Reversing the process just described above, this can be interpreted as a discrete-time representation of the chemical kinetics

$$\begin{aligned} \frac{d[X]}{dt} &= \lambda_X \cdot \frac{[S]}{[S]_{\text{ss}}} \cdot \frac{2}{1+[I]/[I]_{\text{ss}}} - \frac{1}{\tau}[X] \\ &= \lambda_X\, k([I])[S] - \frac{1}{\tau}[X], \qquad\qquad k([I]) = \frac{2}{[S]_{\text{ss}}} \cdot \frac{[I]_{\text{ss}}}{[I]_{\text{ss}} + [I]} \end{aligned}$$

where $[S]_{\text{ss}}$ and $[I]_{\text{ss}}$ describe the steady state concentrations of the stimulus and the inhibitor, respectively. These kinetics can hence be interpreted as first-order synthesis of $X$ due to $S$, with the rate of synthesis being influenced by the concentration of inhibitor in a fashion that is independent of how much stimulant is present. This is in contrast to, say, the kinetics that arise when $S$ and $I$ compete meaningfully over the same receptor.

To complete our justification of the modelling choices made in PSoup, we consider the functional forms that were chosen for the separate types of interaction in turn. We stress, however, that as PSoup is a qualitative modelling approach and its predictions are interpreted in a qualitative fashion, it is not strictly required that a modeller's beliefs about the biological kinetics perfectly match those discussed below. Rather, the arguments we present for deriving the different functional forms used by PSoup serve as evidence of the approach's biochemical backing and highlight the implicit assumptions that are effectively being made when choosing these functional forms.

Firstly, the effects of stimulatory interactions are described in PSoup by

$$f_{i,\text{stim}}(\mathbf{C}) = \frac{\sum_{j \in \mathcal{S}(i)} C_j}{\sum_{j \in \mathcal{S}(i)} 1}$$

where $\mathcal{S}(i)$ denotes the set of nodes that act as stimuli (distinct from *necessary stimuli*, discussed below) of node $i$. The denominator simply counts the number of stimuli to ensure $f_{i,stim}(\mathbf{1}) = 1$, and as the denominator is constant, this function can be understood as separate synthesis of the species $i$ from all of its stimuli. The physical rates of synthesis and steady state concentration of species $i$ are hidden by the non-dimensionalisation and left unspecified in a qualitative PSoup model, but this demonstrates the implicit assumption that under baseline conditions, all of the $i$-th species' stimuli are contributing equally to its synthesis.

The effects of inhibitory interactions are described in PSoup by

$$f_{i,\text{inhib}}(\mathbf{C}) = \frac{1 + \sum_{j \in \mathcal{J}(i)} 1}{1 + \sum_{j \in \mathcal{J}(i)} C_j}$$

where $\mathcal{J}(i)$ describes the set of nodes that inhibit node $i$. This corresponds to all of the inhibitors competing for the same receptor/enzyme, the unavailability of which causes the inhibitory effect (see for example Yonetani and Theorell (1964), in the case where the two inhibitors compete for the same enzyme site and the substrate concentration is treated as fixed). For the purposes of completeness, we present a simpler derivation here. Let us denote the physical concentration of each inhibitor by

$[I_k]$, the free concentration of the receptor/enzyme $[R]$, and its complexes with the inhibitors $[RI_k]$. Then, the total amount of receptor is

$$[R]_{\mathrm{tot}} = [R] + \sum_{k=1}^{N_{\mathrm{inhib}}} [RI_k].$$

For each individual inhibitor, its dynamics are described by

$$\frac{d[I_k]}{dt} = -\alpha_k [R][I_k] + \beta_k [RI_k]$$

and if we assume the binding rates $\alpha_k$ and unbinding rates $\beta_k$ are sufficiently fast, these dynamics can be treated at (quasi) steady state, such that

$$[RI_k] = \frac{\alpha_k}{\beta_k}[R][I_k]$$

Substituting these expressions into the expression for the total concentration of the receptor/enzyme,

$$[R]_{\mathrm{tot}} = [R]\left(1 + \sum_{k=1}^{N_{\mathrm{inhib}}} \frac{\alpha_k}{\beta_k}[I_k]\right)$$

Given this, the ratio of free receptors/enzyme to the total, which defines the reduced rate of activity after taking into account inhibition, is

$$\frac{[R]}{[R]_{\mathrm{tot}}} = \frac{1}{1 + \sum_{k=1}^{N_{inhib}} [I_k]/K_k}, \qquad K_k = \frac{\beta_k}{\alpha_k}.$$

The expression $f_{i,\mathrm{inhib}}(\mathbf{C})$ as used in PSoup is matched to this form by writing

$$f_{i,\mathrm{inhib}}(\mathbf{C}) = \frac{1 + \sum_{j\in\mathcal{J}(i)} 1}{1 + \sum_{j\in\mathcal{J}(i)} C_j} = \frac{1 + \sum_{j\in\mathcal{J}(i)} 1}{1 + \sum_{j\in\mathcal{J}(i)} [C_j]/s_j}$$

and then assuming that the saturation constants for the inhibitors' actions on this node are all given by $K_j = s_j$. Doing so, the denominator precisely matches that for $[R]/[R]_{\mathrm{tot}}$ above, and the numerator is a constant,

$$f_{i,\mathrm{inhib}}(\mathbf{C}) = \frac{1 + \sum_{j\in\mathcal{J}(i)} 1}{1 + \sum_{j\in\mathcal{J}(i)} [C_j]/K_j} = \left(1 + \sum_{j\in\mathcal{J}(i)} 1\right) \cdot \frac{[R]}{[R]_{\mathrm{tot}}}$$

PSoup's implementation of inhibition thus corresponds to reducing the rate of activity due to limited availability of a required enzyme/receptor, that all of the node's inhibitors all together act to occupy. The choice implicitly imposes an assumption that the inhibitors are all equally effective in their inhibitory role under baseline conditions — specifically, each inhibitory species' half-saturation constant is equal to its baseline equilibrium concentration. The maximal rate is given by one plus the total number of inhibitors a node possesses.

Necessary stimuli are treated independently in PSoup, using a product of functions of one variable that represent each of a node's stimuli that are treated as necessary, individually. Analogous to the above, we denote the set of necessary stimuli for the $i$-th node in a PSoup network by $\mathcal{NS}(i)$. Then, the function describing the effect of a node's necessary stimuli in PSoup is given by

$$f_{i,\mathrm{necstim}} = \prod_{j\in\mathcal{NS}(i)} f_{ns}(C_j).$$

The function describing the effect of any one necessary stimulus must satisfy the conditions $f_{ns}(0) = 0$ and $f_{ns}(1) = 1$. This ensures that if any necessary stimulus is absent then the total output is zero, and that the standard baseline condition $f_{i,\mathrm{necstim}}(\mathbf{1}) = 1$. PSoup comes with three different options for this functional form included by default, although the user can also specify their own. The effects of modifiers in PSoup, as used to perturb the values of nodes and represent conditions such as gene knock-out, can also be filtered through the same functional forms used for necessary stimuli. By default, PSoup uses the "linear" function $f(x) = x$, that applies modifiers directly.

Two of these three functional forms were derived by considering a simplified model for the *lac* operon, in which lactose (the necessary stimulus) is required to remove a repressor that blocks a binding site (promoter region) that triggers gene expression. In our simplified description of this system, we model the dynamics of the repressor, $R$, the (necessary) stimulus, referred to as activator $A$, and the binding site $B$. Although in the *lac* operon the repressor binds next to the promoter region and this blocks it, as cells either have the repressor bound or not, and this solely determines the transcription rate in our simplified model, the mathematics are unchanged if the receptor is treated as forming a complex *with* the binding site, and it is the level of free binding site that gives our rate of interest.

We assume no natural unbinding of repressor and binding site, and also that perfect repression of transcription when the repressor is bound, so that the activator is indeed a necessary stimulus for triggering transcription. We also assume that the activator has equal affinity for binding with free repressor, and for cell-bound repressor. Synthesis and turnover are ignored (or treated as at equilibrium), such that the total amounts of activator (free or bound to repressor), repressor (free, bound to activator or bound to the binding site) and binding sites (free, or bound to repressor), are all fixed. We also treat cells as having the repressor bound by default, such that the total number of repressors and binding sites is the same. Mathematically, $[R]_{\text{tot}} = [R]+[BR]+[AR] = [B]+[BR] = [B]_{\text{tot}}$, and $[A]_{\text{tot}} = [A]+[AR]$. Combining these two relationships, $[R] + [BR] = [B]_{\text{tot}} - ([A]_{\text{tot}} - [A])$.

To derive the equilibrium activity rate (the amount of binding sites with no repressor in the way), we begin with the dynamics for the amount of free activator,

$$\frac{d[A]}{dt} = -\alpha_{AR}[A][R] - \alpha_{AR}[A][BR] + \beta_{AR}[AR],$$

where the three terms represent the binding of the activator to free repressor, the binding of the activator to cell-bound repressor, and the disassociation of the activator and the repressor, respectively. Setting the rate of change to zero to determine the proportion of free activator present at equilibrium,

$$\begin{aligned} 0 &= -\alpha_{AR}[A]\left([R] + [BR]\right) + \beta_{AR}[AR] \\ 0 &= -\alpha_{AR}[A]\left([B]_{\text{tot}} - [A]_{\text{tot}} + [A]\right) + \beta_{AR}\left([A]_{\text{tot}} - [A]\right) \\ 0 &= K_{AR}[A]^2 + \left(1 + K_{AR}[B]_{\text{tot}} - K_{AR}[A]_{\text{tot}}\right)[A] - [A]_{\text{tot}}. \end{aligned}$$

Here the second equation is obtained by substituting in the relationships involving total concentrations defined above, and the third by grouping into powers of $[A]$ and defining $K_{AR} = \alpha_{AR}/\beta_{AR}$. Some notable limiting cases here are if activator rapidly unbinds from repressor compared to the rate at which it binds to it (note that a repressor that separates from an activator does not automatically cause it to re-attach to a cell), then $K_{AR} \to 0$ and $[A] \to [A]_{\text{tot}}$. The same limit is arrived at in the case $[A]_{\text{tot}} \gg [B]_{\text{tot}}$ because then there is not enough repressor in the system to meaningfully affect the availability of free $A$.

Next, we consider the dynamics of the concentration of free binding sites,

$$\frac{d[B]}{dt} = -\alpha_{BR}[B][R] + \alpha_{AR}[A][BR],$$

where the two terms represent repressor re-binding to cells, and the binding of the activator to cell-bound repressor (which removes it from blocking the binding site). Again setting the rate of change to zero, and substituting in the relationships in terms of total amounts of chemical species,

$$\begin{aligned} 0 &= -\alpha_{BR}[B][R] + \alpha_{AR}[A][BR] \\ 0 &= -\alpha_{BR}[B]\left([B] - [A]_{\text{tot}} + [A]\right) + \alpha_{AR}[A]\left([B]_{\text{tot}} - [B]\right) \\ 0 &= [B]^2 + \left((1+\kappa)[A] - [A]_{\text{tot}}\right)[B] - \kappa[A][B]_{\text{tot}}, \end{aligned}$$

where $\kappa = \alpha_{AR}/\alpha_{BR}$ denotes the ratio of the repressor's binding rate constant with the activator, and the repressor's binding rate constant with cell binding sites.

We derive functional forms for necessary stimuli in PSoup by considering that the rate of activity will be proportional to the number of free sites, $[B]$, and determining how this varies as a function of the total amount of the activator (necessary stimulus) supplied, $[A]_{\text{tot}}$. This is completely defined by the above two quadratics, with the nature of the functional form set by the values of the constants $K_{AR}$, $\kappa$, and $[B]_{\text{tot}}$. The options already incorporated into PSoup correspond to specific regimes where these dynamics simplify.

The first scenario is when the activator binds much more easily to the repressor than it unbinds from it. In the extreme of this scenario, the equilibrium state has all supplied activator finding as much repressor it can to bind to, hence removing all free activator (if cells are in abundance) or removing all repressor from cells (if activator is in abundance). Mathematically, this scenario corresponds to $K_{AR} \to \infty$, which causes the first quadratic above to simplify to

$$0 = [A]^2 + \left([B]_{\text{tot}} - [A]_{\text{tot}}\right)[A] \quad \Longrightarrow \quad [A] = \{0, [A]_{\text{tot}} - [B]_{\text{tot}}\}$$

The first solution is obtained when the second is physically infeasible (non-negative), and the second is obtained if it is feasible. We then substitute each of these solutions in turn into the second quadratic. Beginning with $[A] = 0$,

$$0 \;=\; [B]\left([B] - [A]_{\text{tot}}\right) \quad \Longrightarrow \quad [B] = \{0, [A]_{\text{tot}}\},$$

and it is the second solution for $[B]$ that corresponds to equilibrium for all species. On the other hand, if we choose the second solution for $[A]$, and substitute $[A] = [A]_{\text{tot}} - [B]_{\text{tot}}$ into the second quadratic, we obtain

$$\begin{aligned} 0 &= [B]^2 + ((1+\kappa)([A]_{\text{tot}} - [B]_{\text{tot}}) - [A]_{\text{tot}})[B] - \kappa[B]_{\text{tot}}([A]_{\text{tot}} - [B]_{\text{tot}}) \\ 0 &= ([B] + \kappa([A]_{\text{tot}} - [B]_{\text{tot}}))([B] - [B]_{\text{tot}}) \qquad \Longrightarrow \qquad [B] = \{\kappa([B]_{\text{tot}} - [A]_{\text{tot}}), [B]_{\text{tot}}\}. \end{aligned}$$

The substitution that leads to this result requires $[A]_{\text{tot}} > [B]_{\text{tot}}$, and given that $\kappa > 0$, the first solution is negative and not physically feasible. Hence, this scenario gives $[B] = [B]_{\text{tot}}$. Summarising the two scenarios together,

$$[B] = \begin{cases} [A]_{\text{tot}} & [A]_{\text{tot}} < [B]_{\text{tot}} \\ [B]_{\text{tot}} & [A]_{\text{tot}} \geq [B]_{\text{tot}} \end{cases}$$

As stated above, the rate of activity (in the *lac* operon case, the rate of gene expression) is what defines the function, $f_{ns}(C)$, and is proportional to the number of free sites, $[B]$. The non-dimensionalised amount of the necessary stimulus species, $C$, is proportional to the physical concentration of activator supplied. However, these proportionality constants are fixed by the baseline condition that $f_{ns}(1) = 1$. Satisfying this condition after substituting $f_{ns}(C) \propto [B]$ and $[A]_{\text{tot}} \propto C$, we may write

$$f_{ns}(C) = \begin{cases} C & C < C_{\max} \text{ and } C_{\max} > 1 \\ C_{\max} & C \geq C_{\max} \text{ and } C_{\max} > 1 \\ C/C_{\max} & C < C_{\max} \text{ and } C_{\max} \leq 1 \\ 1 & C \geq C_{\max} \text{ and } C_{\max} \leq 1. \end{cases}$$

The value of $C_{\max}$ is a free choice (it depends on the value of $[B]_{\text{tot}}$ and on how $C$ is related to $[A]_{\text{tot}}$). In particular, we can obtain the "linear" function $f_{ns}(C) = C$ that is used in PSoup by default for necessary stimuli if $C_{\max} \to \infty$.

The scenario just considered, with very strong binding between the activator and the repressor, allows the supplied amount of activator to very consistently remove the equivalent amount of repressor. In the opposite scenario, the effect is more subtle. When binding between the activator and repressor is weak, they disassociate rapidly, but even a brief binding event removes the repressor from blocking the active region of the cell. Then, because the activator is unbound again, it is free to find another cell-bound repressor and remove it. This results in low amounts of supplied activator actually being *more* effective than in the strong-binding case, but also causes higher amounts of activator to become less effective because it remains ineffective at keeping the repressor occupied and unable to re-bind to cells. It is this scenario that we use to inform the second functional form pre-supplied for use by necessary stimuli in PSoup.

Mathematically, we consider $K_{AR} \to 0$, which reduces the first quadratic to a simple linear equation with solution $[A] = [A]_{\text{tot}}$. This makes sense, as if bound activator disassociates essentially immediately, then all activator supplied will be in the free state. Substituting $[A] = [A]_{\text{tot}}$ into the second quadratic, it simplifies to

$$0 = [B]^2 + \kappa[A]_{\text{tot}}[B] - \kappa[A]_{\text{tot}}[B]_{\text{tot}},$$

which has (feasible) solution

$$[B] = \frac{-\kappa[A]_{\text{tot}} + \sqrt{\kappa^2[A]_{\text{tot}}^2 + 4\kappa[A]_{\text{tot}}}}{2}.$$

This limits to $[B] \to 1$ as $[A]_{\text{tot}} \to \infty$, regardless of the value of $\kappa$. For the form used in PSoup, we choose $C \propto [A]_{\text{tot}}$ to be $C = 2\kappa[A]_{\text{tot}}$, which gives the simple form

$$f_{ns}(C) \propto [B] = \sqrt{C^2 + 2C} - C.$$

Then, choosing the constant of proportionality such that $f_{ns}(1) = 1$,

$$f_{ns}(C) = \frac{\sqrt{C^2 + 2C} - C}{\sqrt{3} - 1}.$$

This is referred to in PSoup as a "switch-like" function, as only a small amount of the stimulus is required to produce a good amount of effect.

The final function pre-supplied for necessary stimuli in PSoup is a simple Michaelis–Menten style relationship. We do not present the derivation for this form here as it is widely available Johnson and Goody (2011), but in short, the reaction velocity $v$ under the Michaelis–Menten kinetics is

$$v = \frac{v_{\max}[S]}{K + [S]}.$$

Here $[S]$ is the concentration of substrate, $K$ is the ratio of binding and un-binding between enzyme and substrate, and $v_{\max}$ is the maximum velocity for the reaction. For PSoup, the substrate takes on the role of the necessary stimulus, and we choose $K = 1$ for simplicity and $v_{\max} = 2$ such that the resultant function satisfies the baseline condition, $f_{ns}(1) = 1$. These choices result in

$$f_{ns}(C) = \frac{2C}{1 + C}.$$

### Supplementary Seven

This supplement contains the full comparison of simulated conditions for each of the three alternative networks explored. Here, only one representative for each condition is shown, excluding all baseline conditions given that they by definition will be correct.

Figure 15 shows the comparison between the biological results and simulated outcomes for the alternative network when the stimulatory edge between sucrose and cytokinin has been removed. This network had an accuracy of 81.6%.

Figure 16 shows the comparison between the biological results and simulated outcomes for the alternative network when an additional edge between sucrose and the branching node had been added. This network had an accuracy of 86.8%.

Figure 17 shows the comparison between the biological results and simulated outcomes for the alternative network when the stimulatory edge between sucrose and cytokinin has been removed, and an additional edge between sucrose and the branching node had been added. This network had an accuracy of 84.2%.

### Supplementary Eight

When building a model based on a diagrammatic input constructed with SBGN and the Newt editor, PSoup will create a folder to store all the components that make up that model. There are several files that are automatically generated by PSoup. The most important is the nextStep.R script, which contains an R function that creates and defines all the equations for calculating the values of nodes at each timestep. When running simulations, this is the function that will be used to iterate the model. By producing this script, PSoup has made the generated model completely transparent to the user.

In addition to this script, several supporting files are produced defining the simulations to be run. These definition files are used to specify the starting values of nodes at the beginning of the simulation (nodeStartDef.RData), and to specify the weighting of any node that is to be modified (modifierDef.RData). Given that PSoup needs to be able to run simulations on whatever network that is provided to it, the nodeStartDef object becomes important for two separate functions. First, it defines the internal data structure that is used to record the progression of the simulation. Second, it provides the actual starting values for the simulation, which will be progressively updated using the update rules defined in the nextStep function. While the nodes themselves are variable, those of the modifiers represent non-changing inputs to the simulations.

To define a set of simulations, users can edit either of the definition files. For each file, each row represents one simulation. Once these files have been defined, they are saved back into the model definition folder. In this way, to run the set of simulations, all the user needs to do is provide the path directory of the model file to the setupSims function. PSoup will automatically cycle through rows in the definition files to produce an output object. The output file produced by setupSims, will be saved directly to the model output folder provided as input to the simulation. For examples on how to set up such a simulation, refer to the PSoup tutorials at `https://nicolezfortuna.github.io/PSoup/`.

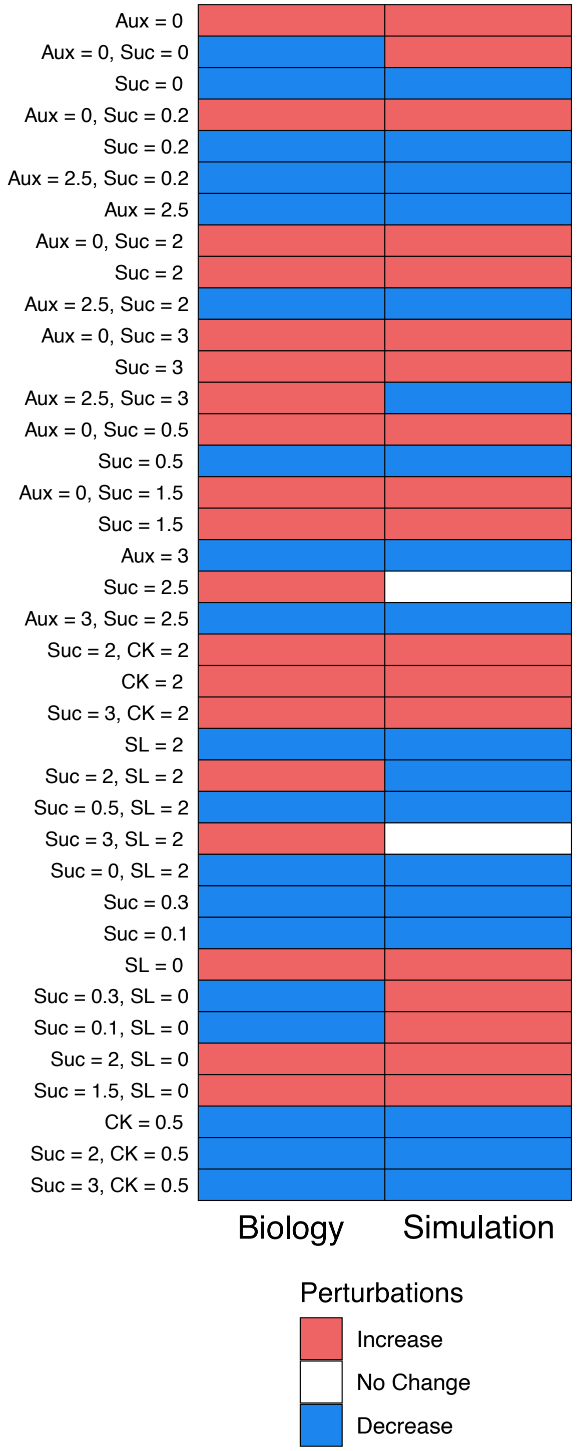


**Figure 15** Comparing the accuracy of the alternative network where the edge between sucrose and cytokinin has been removed. Here, only unique conditions have been represented. On the left are the biological outcomes, on the rights are the simulated outcomes. On the y-axis are the conditions for nodes relative to the baseline.

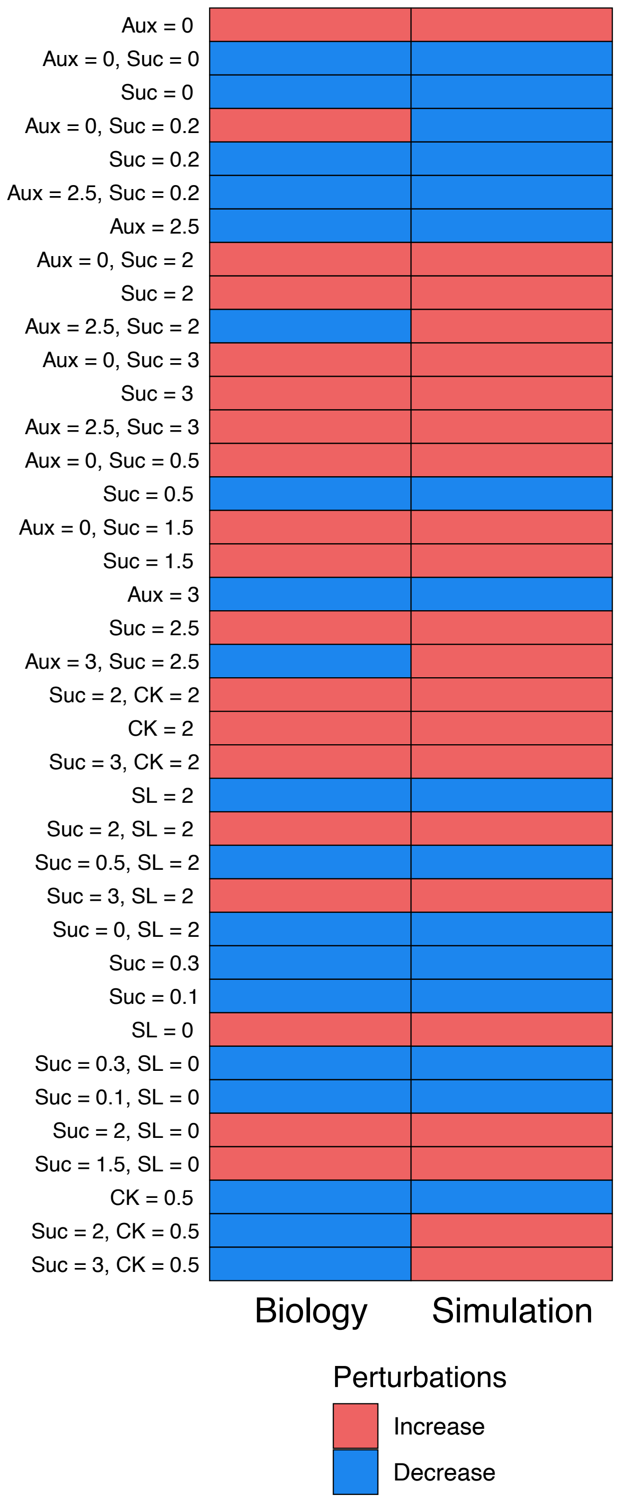


**Figure 16** Comparing the accuracy of the alternative network where there is an additional edge between sucrose and the branching node. Here, only unique conditions have been represented. On the left are the biological outcomes, on the rights are the simulated outcomes. On the y-axis are the conditions for nodes relative to the baseline.

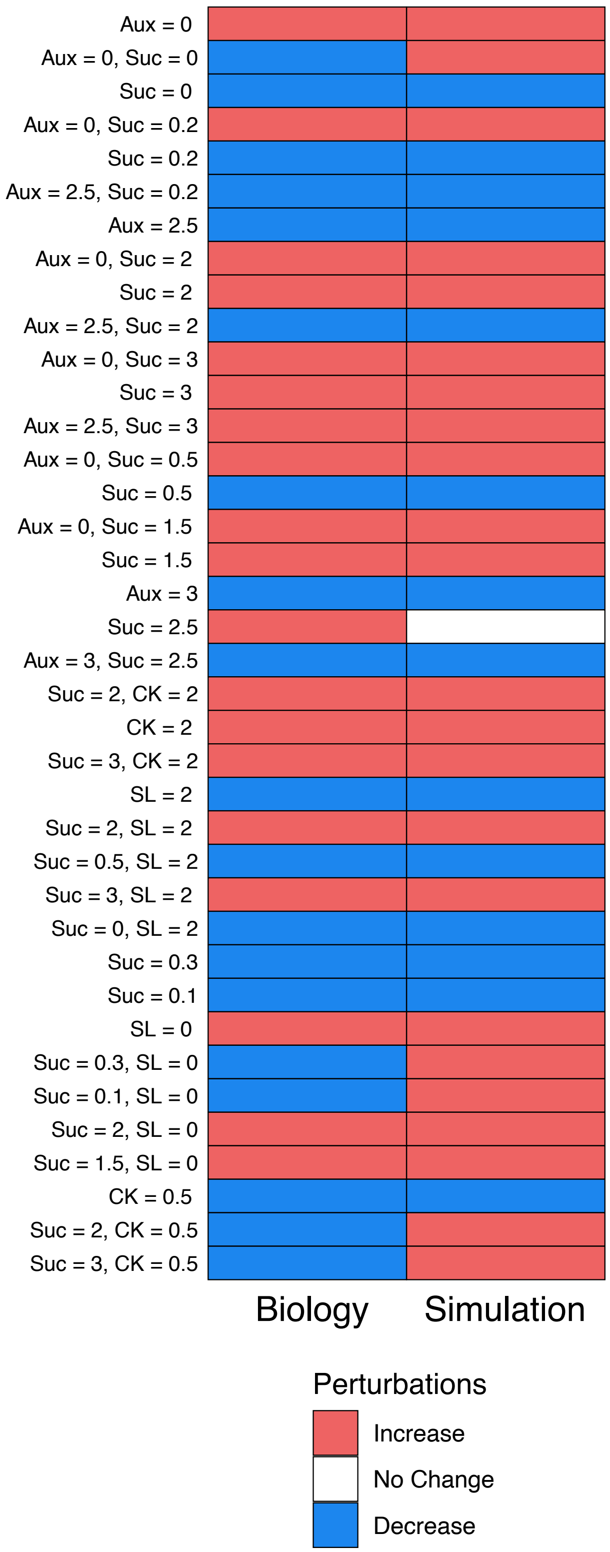


**Figure 17** Comparing the accuracy of the alternative network where the edge between sucrose and cytokinin has been removed and there is an additional edge between sucrose and the branching node. Here, only unique conditions have been represented. On the left are the biological outcomes, on the rights are the simulated outcomes. On the y-axis are the conditions for nodes relative to the baseline.

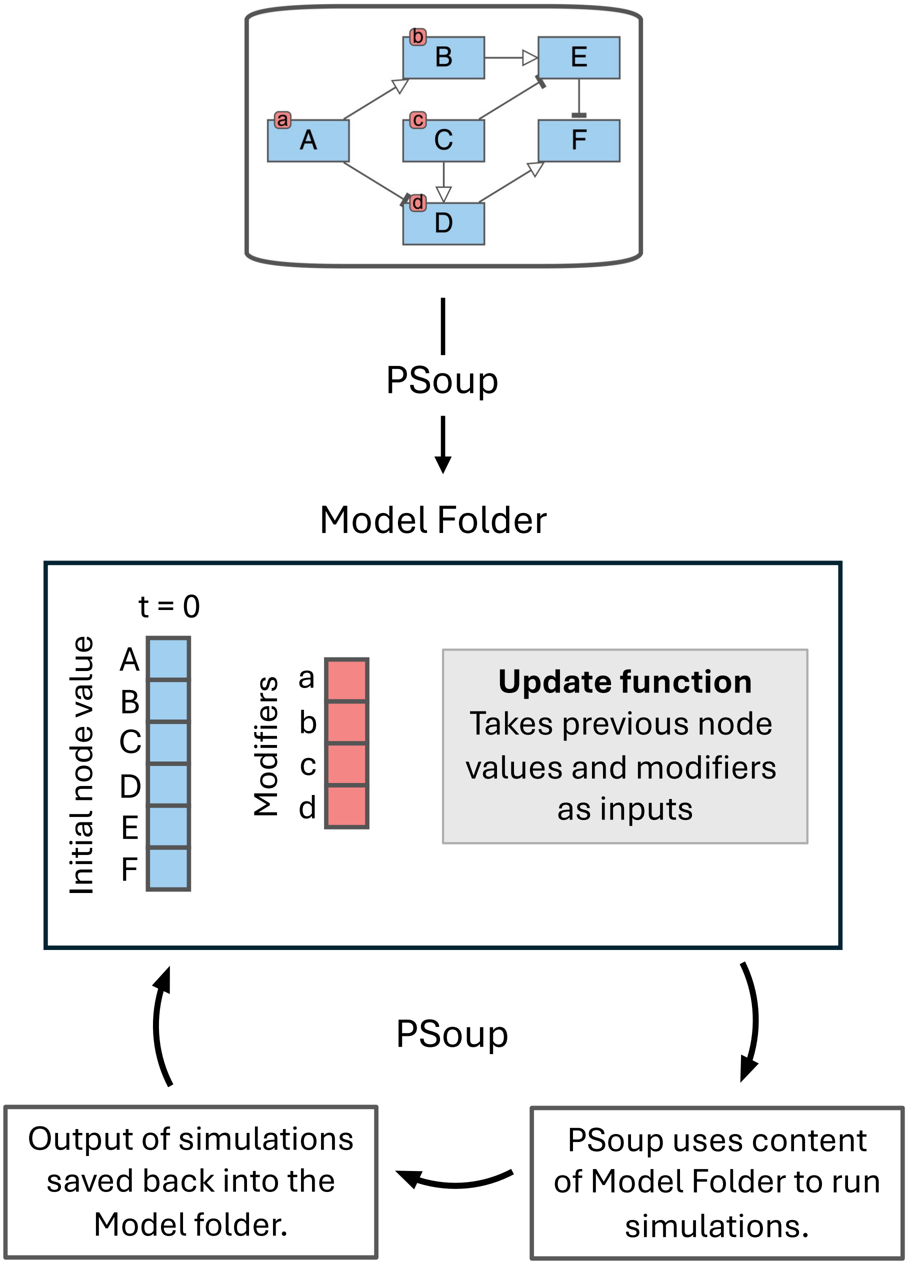


**Figure 18** Flow chart depicting the creation of a Folder defining the model using PSoup. PSoup will then use the contents of the Model Folder to run simulations. The output of these simulations is then saved back into the Model Folder.
